# A generalized cortical activity pattern at internally-generated mental context boundaries during unguided narrative recall

**DOI:** 10.1101/2021.09.07.459300

**Authors:** Hongmi Lee, Janice Chen

**Affiliations:** Department of Psychological and Brain Sciences, Johns Hopkins University, Baltimore, MD 21218, USA

## Abstract

Current theory and empirical studies suggest that humans segment continuous experiences into events based on the mismatch between predicted and actual sensory inputs; detection of these “event boundaries” evokes transient neural responses. However, boundaries can also occur at transitions between internal mental states, without relevant external input changes. To what extent do such “internal boundaries” share neural response properties with externally-driven boundaries? We conducted an fMRI experiment where subjects watched a series of short movies and then verbally recalled the movies, unprompted, in the order of their choosing. During recall, transitions between movies thus constituted major boundaries between internal mental contexts, generated purely by subjects’ unguided thoughts. Following the offset of each recalled movie, we observed stereotyped spatial activation patterns in the default mode network, especially the posterior medial cortex, consistent across different movie contents and even across the different tasks of movie watching and recall. Surprisingly, the between-movie boundary patterns did not resemble patterns at boundaries between events within a movie. Thus, major transitions between mental contexts elicit neural phenomena shared across internal and external modes and distinct from within-context event boundary detection, potentially reflecting a cognitive state related to the flushing and reconfiguration of situation models.

## Introduction

Humans perceive and remember continuous experiences as discrete events (Brunec et al., 2018; Clewett et al., 2019; Shin & DuBrow, 2021; Zacks, 2020). Studies of event segmentation have shown that when participants attend to external information (e.g., watch a video), 1) boundaries between events are detected when mismatches arise between predicted and actual sensory input (Zacks et al., 2007, 2011), and 2) boundary detection evokes transient neural responses in a consistent set of brain areas (Reagh et al., 2020; Speer et al., 2007; Zacks et al., 2001). Among these areas is the default mode network (DMN; Buckner & DiNicola, 2019) proposed to be involved in representing complex mental models of events (Ranganath & Ritchey, 2012; Ritchey & Cooper, 2020). However, a substantial portion of human cognition is internally-driven (Hasselmo, 1995; Honey et al., 2017), and such spontaneous production of thoughts and actions is also punctuated by mental context transitions (Christoff et al., 2016; Mildner & Tamir, 2019; Smallwood & Schooler, 2015; Tseng & Poppenk, 2020). What manner of brain activity marks boundaries between mental contexts when they are internally-generated? Are the brain responses at internal boundaries similar to those at external boundaries?

Here, we used naturalistic movie viewing and free spoken recall with fMRI to characterize neural activity at boundaries between internally-generated mental contexts (Figure 1A). Subjects watched ten short movies (encoding phase), then verbally recounted the movies in any order, in their own words (recall phase). The transitions between recalled movies were determined purely by subjects’ internal mentation; no external cues prompted the recall onset or offset of each movie. Moreover, the unguided spoken recall allowed us to identify the exact moments of context transitions and explicitly track shifts in the contents of thoughts (Chen et al., 2017; Sripada & Taxali, 2020), which was not possible in prior studies using silent rest (Karapanagiotidis et al., 2020; Tseng & Poppenk, 2020). At these internal boundaries between recalled movies, we observed transient, highly generalizable and fine-grained activation patterns throughout the DMN, consistent across diverse movie contents and similar to those at external between-movie boundaries during encoding. Moreover, these between-movie boundary patterns were not merely stronger versions of within-movie “event boundaries”, but instead manifested as a distinct type of neural transition. We propose that these cortical patterns reflect a cognitive state related to the major flushing and reconfiguration of mental context (DuBrow et al., 2017; Manning et al., 2016).

**Figure 1.**
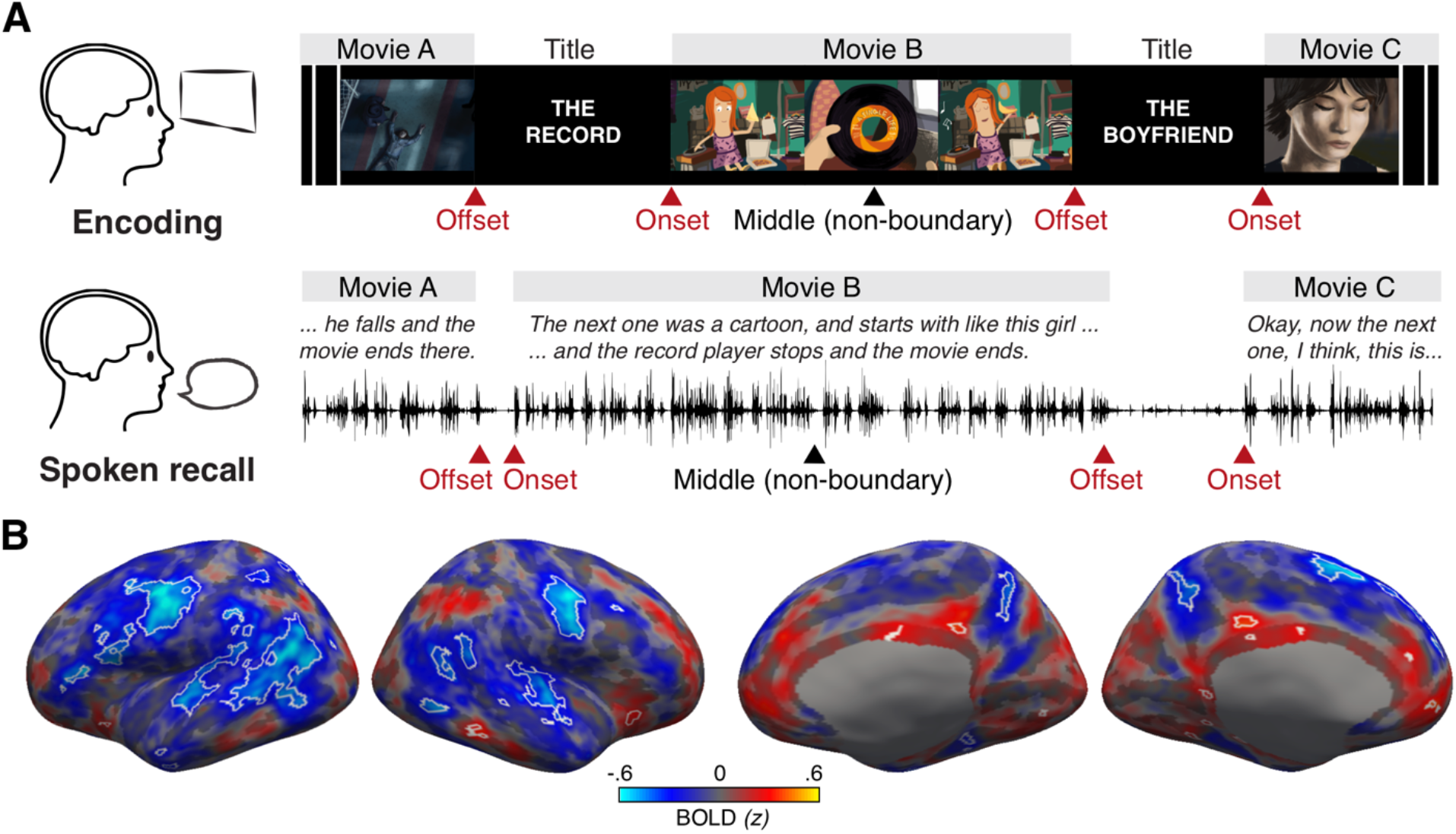
Experimental procedures and univariate responses. (A) In the encoding phase, subjects watched 10 short movies approximately 2 to 8 minutes long. Each movie started with a 6-second title scene. In the free spoken recall phase, subjects verbally recounted each movie plot in as much detail as possible regardless of the order of presentation. After recalling one movie, subjects spontaneously proceeded to the next movie, and the transitions between movies were considered as internally-driven boundaries. Red arrows indicate the boundaries (onsets and offsets) between watched or recalled movies. Black arrows indicate the non-boundary moments (middle) of each watched or recalled movie. (B) Whole-brain maps of unthresholded mean activation (BOLD signals z-scored across all volumes within a scanning run) following between-movie boundaries during recall (4.5 – 19.5 seconds from the offset of each movie). Blue areas indicate regions with lower-than-average activation, where the average activation of a scanning run was z = 0. Likewise, red areas indicate regions with higher-than-average activation. White outlines indicate areas that showed significantly lower or higher activation following between-movie boundaries compared to non-boundary periods (FDR corrected *q* < .05; minimum surface area = 16 mm^2^). The non-boundary periods were defined as the middle 15 seconds of each recalled movie, shifted forward by 4.5 seconds. Changes in whole-brain univariate responses across time around the boundaries are shown in Figure 1—figure supplement 1 (recall phase) and Figure 1—figure supplement 2 (encoding phase).

## Results

We first examined whether internally-driven boundaries evoke changes in blood oxygen level-dependent (BOLD) signals during recall. We observed transient changes in activation at the boundaries between recalled movies in widespread cortical regions (Figure 1—figure supplement 1; see Figure 1—figure supplement 3 for activation time courses). A whole-brain analysis with multiple comparisons correction revealed that the mean activation of boundary periods (15 seconds following the offset of each movie) was generally lower than that of non-boundary periods (middle 15 seconds within each movie) in multiple areas including the motor, auditory, and inferior parietal cortices, although a smaller number of regions showed higher activation during non-boundary periods (Figure 1B).

Next, we tested whether there were neural activation patterns specific to internally-driven boundaries and consistent across different movies. We performed a whole-brain pattern similarity analysis on the recall data to identify regions where 1) boundary period activation patterns were positively correlated across different recalled movies (Figure 2A, blue arrow *a* > 0), and 2) this correlation was higher at boundaries than at non-boundaries (Figure 2A, blue arrows *a* > *b*). We observed a consistent boundary pattern, i.e., whenever participants transitioned from talking about one movie to the next, in several cortical parcels (Schaefer et al., 2018) including the DMN and auditory/motor areas (Figure 2B). Thus, the boundary patterns within the recall phase were likely to be driven by both shared low-level sensory/motor factors (e.g., breaks in recall speech generation) as well as cognitive states (e.g., memory retrieval) at recall boundaries. No cortical parcel showed significantly negative correlations between boundary patterns or greater correlations in the non-boundary compared to boundary conditions.

**Figure 2.**
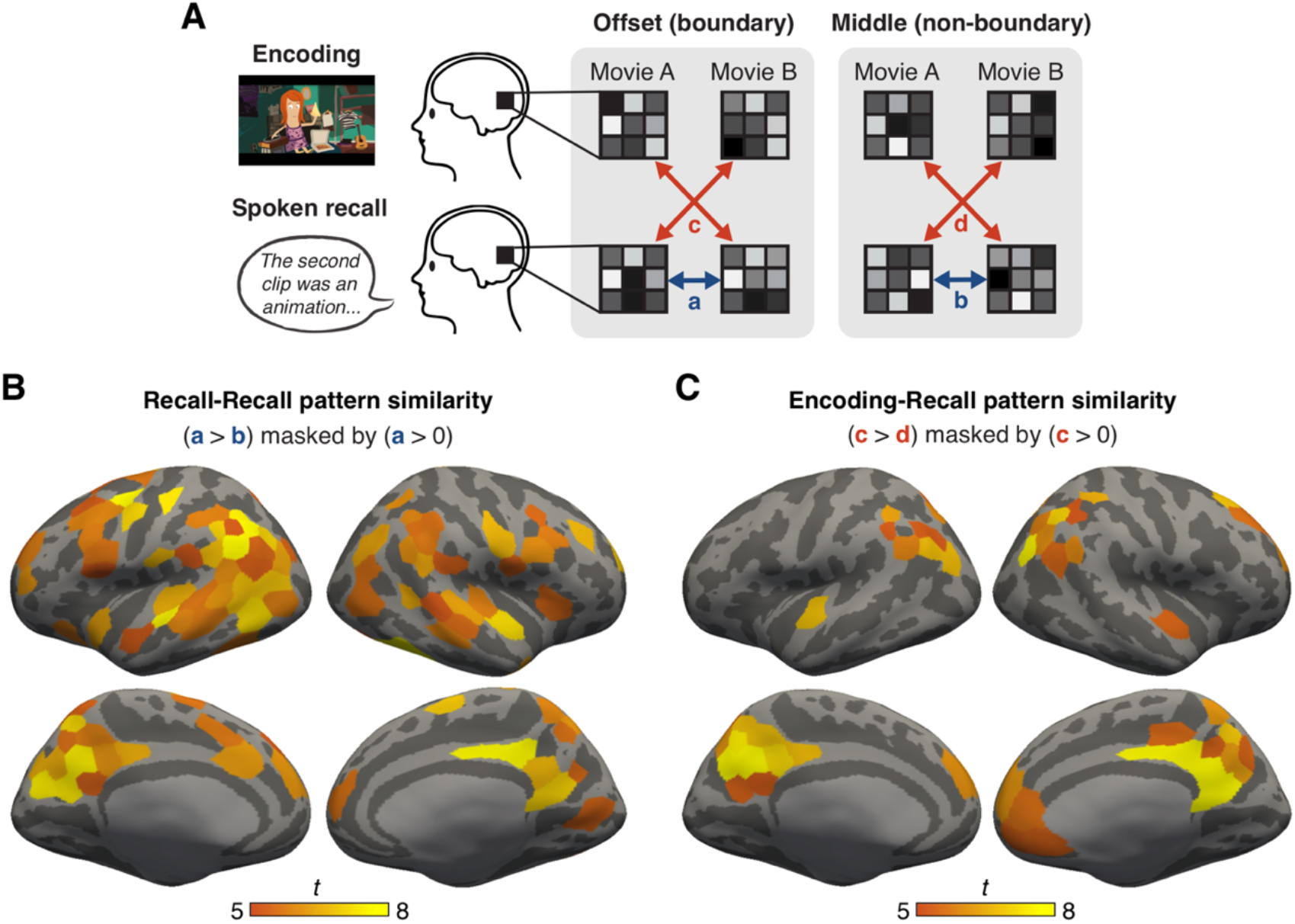
Consistent activation patterns associated with between-movie boundaries. (A) Schematic of the pattern similarity analysis. Boundary patterns were defined as the mean pattern averaged across 15 seconds following the offset of each watched or recalled movie. Non-boundary patterns were defined as the mean pattern averaged across 15 seconds in the middle of each watched or recalled movie. For each subject and cortical parcel (Schaefer et al., 2018; 200 parcels per hemisphere), we computed pairwise between-movie pattern similarity (Pearson correlation), separately for boundary patterns and non-boundary patterns measured during recall (a & b, blue arrows). We also computed between-movie and between-phase (encoding-recall) pattern similarity, again separately for boundary and non-boundary patterns (c & d, red arrows). The time windows for both boundary and non-boundary periods were shifted forward by 4.5 seconds to account for the hemodynamic response delay. (B) Whole-brain *t* statistic map of cortical parcels that showed consistent between-movie boundary patterns during recall. These parcels displayed significantly greater between-movie pattern similarity in the boundary condition compared to the non-boundary condition during recall. The map was masked by parcels that showed significantly positive between-movie pattern similarity in the boundary condition during recall. Both effects were Bonferroni corrected across parcels (*p* < .05). (C) Whole-brain *t* statistic map of cortical parcels that showed consistent between-movie boundary patterns across encoding and recall. These parcels displayed significantly greater between-movie and between-phase pattern similarity in the boundary condition compared to the non-boundary condition. The map was masked by parcels that showed significantly positive between-movie and between-phase pattern similarity in the boundary condition. Both effects were Bonferroni corrected across parcels (*p* < .05).

To what extent is the internally-driven boundary pattern, measured during recall, similar to patterns observed at boundaries during encoding? To test this, we again computed between-movie pattern similarity for all cortical parcels in the brain, but now across the encoding and recall phases (Figure 2A, red arrows). We found that DMN areas showed a consistent boundary pattern across task phases (encoding and recall) and across movies (Figure 2C). Again, no cortical parcel showed negative correlations between boundary patterns or greater correlations in the non-boundary condition. Among the DMN areas, the posterior medial cortex (PMC) showed the most consistent boundary patterns; thus, we next examined the phenomenon in more detail specifically in PMC. Figures 3A and 3C visualize the high and consistently positive correlations of PMC boundary patterns across different movies both within the recall phase (Recall Offset vs. Recall Offset, *t*(14) = 11.82, *p* < .001, Cohen’s *d*_z_ = 3.05, 95% confidence interval (CI) = [.28, .41]) and even between experimental phases (Recall Offset vs. Encoding Offset, *t*(14) = 14.54, *p* < .001, Cohen’s *d*_z_ = 3.75, 95% CI = [.28, .38]). No such correlation was present between non-boundary patterns (*t*(14)s < 1, *p*s > .3). Individual subjects’ activation maps visualize the similarity between boundary patterns during encoding and recall (Figure 3B, Figure 3—figure supplement 1). We observed similar results in the lateral parietal DMN sub-region (angular gyrus; Figure 3—figure supplement 2), as well as using shorter (4.5 s) time windows of boundary and non-boundary periods (Figure 2—figure supplement 1, Figure 3— figure supplement 3).

**Figure 3.**
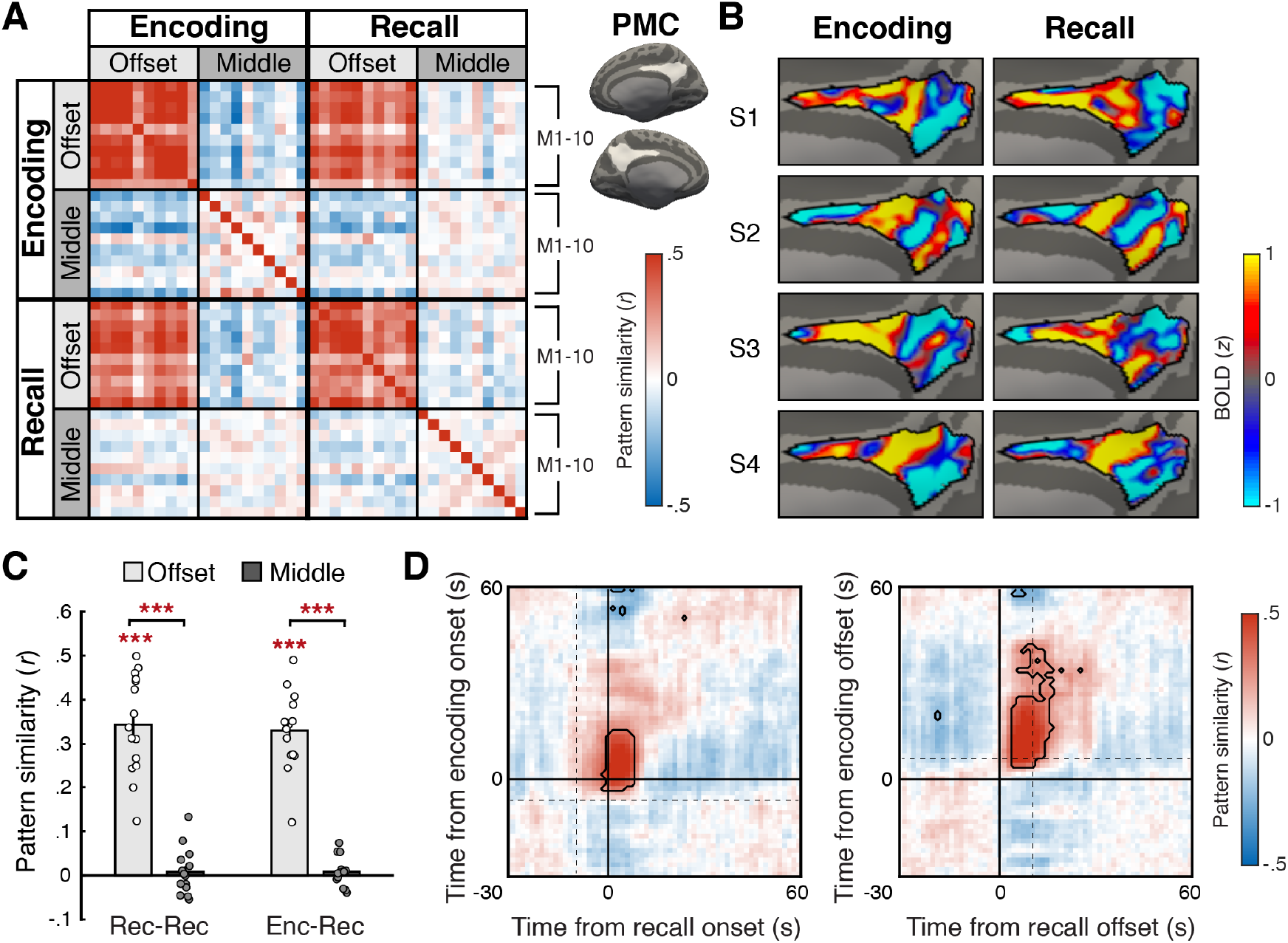
Boundary pattern in the posterior medial cortex (PMC). (A) PMC activation pattern similarity (Pearson correlation) between the 10 movie stimuli (M1 – 10), conditions (Offset = boundary, Middle = non-boundary), and experimental phases (Encoding, Recall), averaged across all subjects. The boundary pattern of a movie was defined as the mean pattern averaged across the 15-second window following the offset of the movie. The non-boundary pattern was defined as the mean pattern averaged across the 15-second window in the middle of a movie. The time windows for both boundary and non-boundary patterns were shifted forward by 4.5 seconds to account for the hemodynamic response delay. PMC regions of interest are shown as white areas on the inflated surface of a template brain. (B) Subject-specific mean activation patterns associated with between-movie boundaries during encoding (left) and recall (right). The boundary patterns were averaged across all movies and then *z*-scored across vertices within the PMC ROI mask, separately for each experimental phase. PMC (demarcated by black outlines) of four example subjects (S1 – 4) are shown on the medial surface of the right hemisphere of the fsaverage6 template brain. (C) Within-phase (Recall-Recall) and between-phase (Encoding-Recall) pattern similarity across different movies, computed separately for the boundary (Offset) and non-boundary (Middle) patterns in PMC. Bar graphs show the mean across subjects. Circles represent individual subjects. Error bars show SEM across subjects. \*\*\**p* < .001. (D) Time-point-by-time-point PMC pattern similarity across the encoding phase and recall phase activation patterns around between-movie boundaries, averaged across all subjects. The time series of activation patterns were locked to either the onset (left) or the offset (right) of each movie. During encoding, the onset of a movie and the offset of the preceding movie were separated by a 6-second title scene. During recall, onsets and offsets of recalled movies were separated by, on average, a 9.3-second pause (boundaries concatenated across subjects, SD = 16.8 seconds). Dotted lines on the left and right panels indicate the mean offset times of the preceding movies and the mean onset times of the following movies, respectively. Note that in this figure, zero corresponds to the true stimulus/behavior time, with no shifting for hemodynamic response delay. Areas outlined by black lines indicate correlations significantly different from zero after multiple comparisons correction (Bonferroni corrected *p* < .05). Time-time correlations within each experimental phase can be found in Figure 3—figure supplement 4.

Thus far, we tested boundary responses following offsets, based on prior findings that post-stimulus neural responses contribute to memory formation (Ben-Yakov et al., 2013; Ben-Yakov & Dudai, 2011; Medvedeva et al., 2021). However, other studies also reported neural responses specific to the onset of an episode (Bulkin et al., 2020; Fox et al., 2005; Wen et al., 2020). Is the generalized boundary pattern evoked by the onset of a movie, rather than the offset? We examined this question by comparing the temporal emergence of the generalized boundary pattern following movie offsets versus onsets (Figure 3D); note that the offset of a movie was temporally separated from the onset of the following movie during both encoding and recall (see Figure 1A). Specifically, we extracted the mean time series of PMC activation patterns around between-movie boundaries, time-locked to either the onset or offset of each watched or recalled movie. We then computed between-phase (encoding-recall) pattern similarity across the individual time points of the activation pattern time series. We found that significantly positive between-phase correlations emerged well before the encoding and recall onsets (Figure 3D, left panel), starting from 4.5 seconds following the offsets of the preceding watched or recalled movie (Figure 3D, right panel). Thus, boundary patterns were not exclusively triggered by movie onsets; it is likely that offset responses significantly contributed to the boundary patterns.

We focused our analyses up to this point on transitions between movies because they provided clear boundaries between mental contexts during recall. However, event boundaries in naturalistic movie stimuli are often defined as transitions between scenes within a movie (Baldassano et al., 2017; Chen et al., 2017; Zacks et al., 2010). In prior work, it has been shown that for within-movie event boundaries, neural responses scale positively with human judgments of the “strength” of scene transitions (Ben-Yakov & Henson, 2018). Thus, we hypothesized that boundaries between movies (i.e., between mental contexts) would manifest as stronger versions of within-movie boundaries with qualitatively similar patterns; in other words, boundary patterns would generalize across different scales of boundaries. To test this idea, we first confirmed that there were consistent within-movie event boundary patterns in PMC during encoding; within-movie boundary patterns were more similar to each other than to non-boundary patterns (Figure 4—figure supplement 1). We then tested whether this within-movie boundary pattern resembled the between-movie boundary pattern, by measuring the correlation between 1) the mean between-movie boundary pattern during recall and 2) the mean within-movie event boundary pattern during encoding (Figure 4). Surprisingly, the two were negatively correlated (*t*(14) = 5.10, *p* < .001, Cohen’s *d*_z_ = 1.32, 95% CI = [-.34, -.14]), in contrast to the strong positive correlation across encoding and recall between-movie boundary patterns (*t*(14) = 25.02, *p* < .001, Cohen’s *d*_z_ = 6.46, 95% CI = [.67, .79]). The within-movie event boundary pattern was also negatively correlated with the encoding phase between-movie boundary pattern (*t*(14) = 7.31, *p* < .001, Cohen’s *d*_z_ = 1.89, 95% CI = [-.44, -.24]). Within-movie and between-movie boundary patterns did not resemble each other, regardless of the specific time windows used to define the boundary periods (Figure 4—figure supplement 2). These results suggest that the between-movie boundary pattern may reflect a cognitive state qualitatively different from the state elicited by within-movie event boundaries during movie watching.

**Figure 4.**
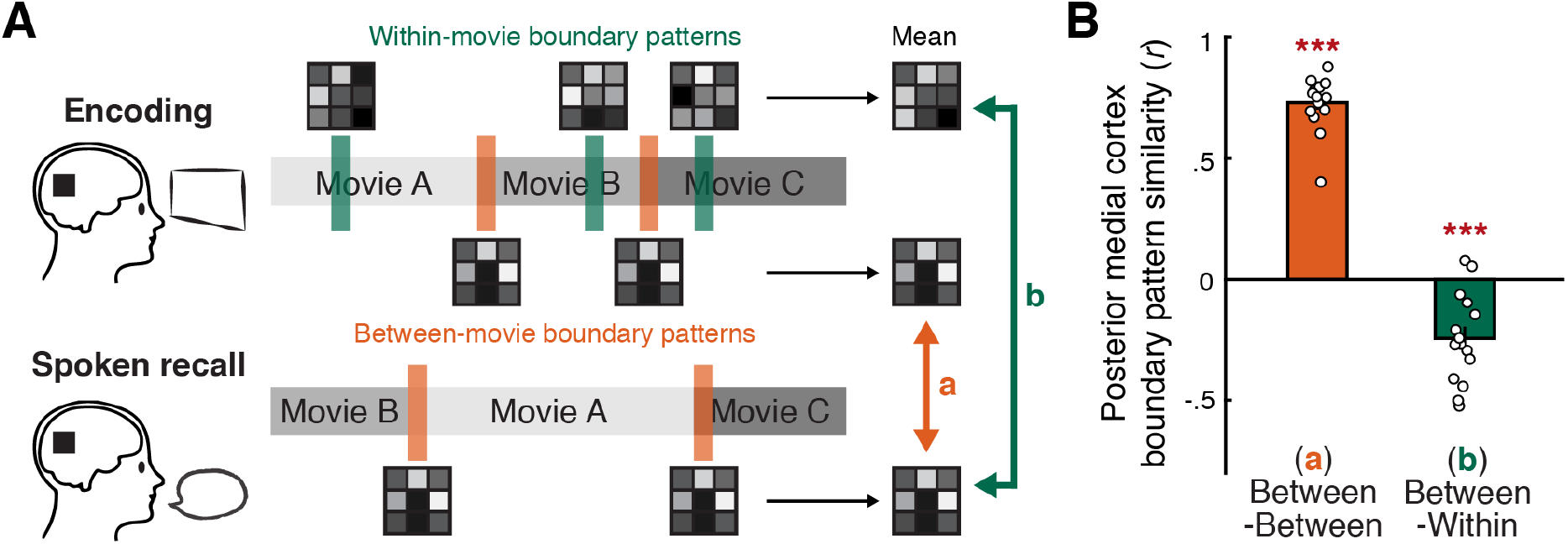
Comparing between-movie and within-movie boundary patterns in the posterior medial cortex (PMC). (A) Schematic of the analysis. For each subject, we created the template PMC activation pattern associated with between-movie boundaries by averaging activation patterns following the offset of each between-movie boundary (orange bars), separately for encoding and recall phases. Likewise, the template within-movie event boundary pattern was created by averaging the activation patterns following the offset of each within-movie boundary during encoding (green bars). We then measured the similarity (Pearson correlation) between the mean between-movie boundary patterns during encoding and recall (a, orange arrow). We also measured the similarity between the mean within-movie boundary pattern during encoding and the mean between-movie boundary pattern during recall (b, green arrow). For both between- and within-movie boundaries, boundary periods were 15-seconds long, shifted forward by 4.5 seconds. (B) Pattern similarity between template boundary patterns. The orange bar shows the mean correlation across the between-movie boundary patterns during encoding and recall. The green bar shows the mean correlation across the between-movie boundary pattern during recall and the within-movie boundary pattern during encoding. Circles represent individual subjects. Error bars show SEM across subjects. \*\*\**p* < .001 against zero.

Is the generalized between-movie boundary pattern driven by shared low-level perceptual or motoric factors rather than cognitive states? First, shared visual features at between-movie boundaries (i.e., black screen) cannot explain the transient, boundary-specific similarity between encoding and recall phases, because visual input was identical across boundary and non-boundary periods during recall (i.e., a fixation dot on black background). Indeed, encoding boundary patterns were more similar to recall boundary patterns than to recall non-boundary patterns in DMN areas, suggesting a limited contribution of shared visual input to the generalized boundary pattern (Figure 2—figure supplement 2). Likewise, the absence of verbal responses at boundaries cannot explain the boundary pattern generalized across encoding and recall phases, as no speech was generated throughout the entire encoding phase. Moreover, PMC boundary patterns showed positive between-phase pattern correlations (*t*(14) = 3.94, *p* = .003, Cohen’s *d*_z_ = 1.25, 95% CI = [.1, .36]) greater than those of non-boundary patterns (*t*(14) = 3.22, *p* = .011, Cohen’s *d*_z_ = 1.02, 95% CI of the difference = [.06, .36]) even when restricted to boundaries without pauses between recalled movies. We also ruled out the possibility that silence during movie title scenes and pauses at recall boundaries drove the generalized boundary pattern in PMC; the recall boundary pattern was not correlated with the pattern associated with silent periods during encoding (*t*(14) = 1.93, *p* = .074, Cohen’s *d*_z_ = .498, 95% CI = [-.19, .01]), whereas the auditory cortex showed a positive correlation between the two (*t*(14) = 10.31, *p* < .001, Cohen’s *d*_z_ = 2.66, 95% CI = [.3, .45]) (Figure 5). Likewise, the movies’ audio amplitudes modulated the time course of similarity between the recall boundary pattern and the encoding data in the auditory cortex, but not in PMC (Figure 5— figure supplements 1, 2).

**Figure 5.**
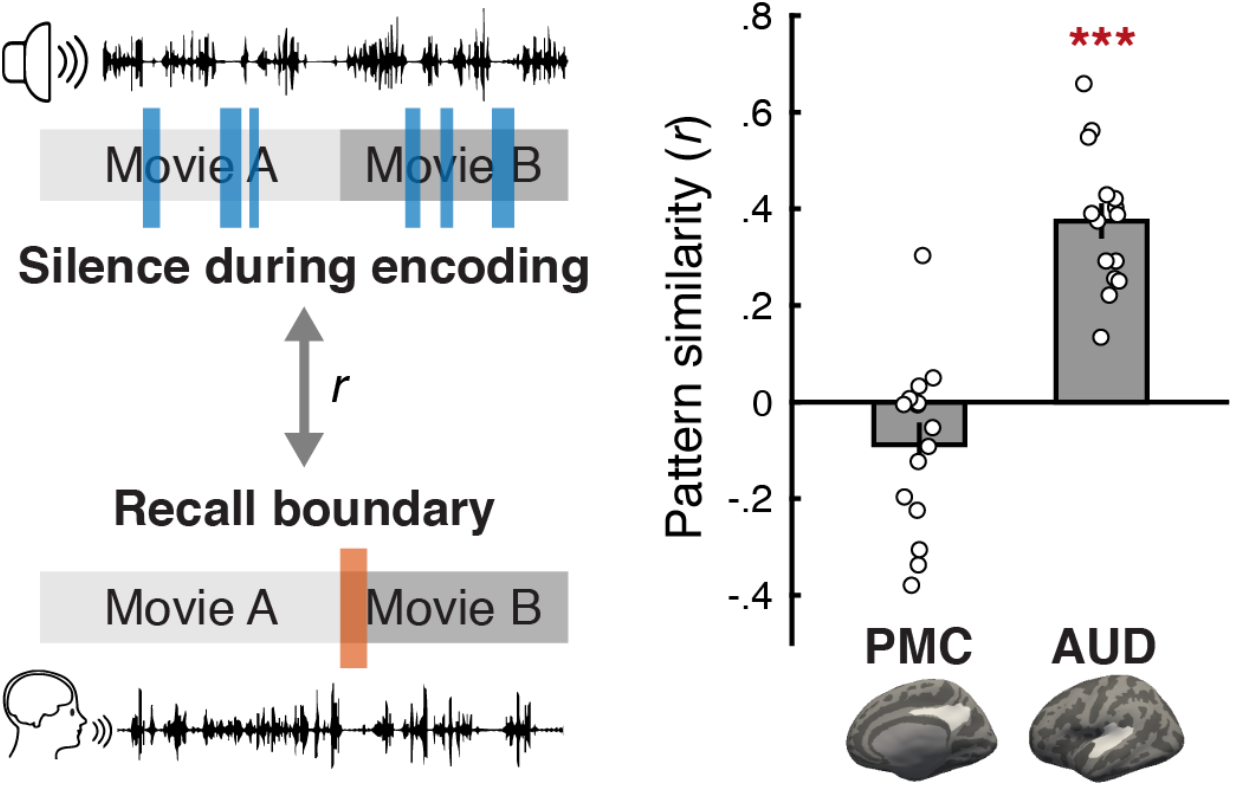
Examining the effects of silence on the generalized boundary pattern. For each subject, we computed a Pearson correlation between the mean activation pattern of the moments of silence during encoding (blue bars) and the mean activation pattern of between-movie boundaries during recall (orange bar) in the posterior medial cortex (PMC) and the auditory cortex (AUD). The moments of silence near between-movie boundaries (i.e., within the first 45 seconds of each movie) during encoding were excluded from the analysis. PMC and AUD regions of interest are shown as white areas on the inflated surface of template brains. Gray bars on the right panel indicate the mean pattern similarity across subjects. Circles represent individual subjects. Error bars show SEM across subjects. \*\*\**p* < .001 against zero.

## Discussion

The current study investigated brain responses to internally-generated boundaries between mental contexts during continuous and unguided memory recall of naturalistic narratives. We found that internally-driven mental context boundaries evoke generalized neural activation patterns in core posterior-medial areas of the DMN (Ritchey & Cooper, 2020). These cortical patterns were similar to those observed at major boundaries between externally-driven contexts (different audiovisual movies), suggesting that they reflect a general cognitive state associated with mental context transitions. However, these between-context patterns were distinct from within-context event boundary detection signals.

The highly similar neural activation patterns for internally- and externally-driven boundaries observed in this study demonstrate event segmentation without changes in external input. This finding diverges from the currently dominant empirical and theoretical perspectives on event segmentation; in most studies, event boundaries are defined or manipulated by changes in perceptual or spatiotemporal features (e.g., Chen et al., 2017; DuBrow & Davachi, 2013; Radvansky & Copeland, 2006), and boundary detection is posited to occur when those changes mismatch our expectations of the current situation (Zacks et al., 2007, 2011). This prediction error framework successfully explains various phenomena related to event perception and memory organization (see Zacks, 2020 for a review); however, evidence has also shown that predicted changes in external features can create boundaries and have similar behavioral effects (Pettijohn & Radvansky, 2016; Schapiro et al., 2013). To resolve the discrepancy, an alternative theoretical framework has recently proposed that boundaries are perceived when the probability distribution of inferred current situations, rather than observed external features per se, changes from the previous time point (Shin & DuBrow, 2021). According to this account, event segmentation can occur when there is no perceptual change or when transitions are already predicted, which may explain the boundary-related neural responses at self-generated transitions between memories during recall in our study.

The boundary pattern which generalized across internally- and externally-driven boundaries was most strongly observed in the DMN, in line with earlier findings implicating the DMN in mental context transitions (Baldassano et al., 2017; Crittenden et al., 2015; Smith et al., 2018). Prior studies have shown that the DMN responds to external context transitions including experimental task switching (Crittenden et al., 2015; Smith et al., 2018) as well as event boundaries in movie clips (Reagh et al., 2020; Speer et al., 2007). Considering these findings and the widely-known involvement of the DMN in internally-oriented cognition (e.g., Addis et al., 2007; Andrews-Hanna et al., 2010; Christoff et al., 2009) together, it has been suggested that the DMN integrates both internal and external information to represent and maintain an abstract mental model of the current situation or state (Stawarczyk et al., 2021; Yeshurun et al., 2021); located furthest away from sensory-motor areas (Smallwood et al., 2021), the DMN integrates information across different modalities (Bonnici et al., 2016; Ramanan et al., 2018) and over long timescales (Chang et al., 2021; Hasson et al., 2015). Supporting this idea, neural activation patterns in sub-regions of the DMN, especially PMC, tend to persist for extended periods of time during naturalistic movie watching, and transitions between these persistent neural states coincide with perceived event boundaries (Baldassano et al., 2017; Geerligs et al., 2021). Our study extends this finding by identifying a transient, boundary-induced phenomenon which is a unique and independent state represented in the DMN. That is, at major event boundaries, a temporary boundary state may exist in between the neural patterns representing the two events, rather than one event pattern switching directly to the next.

Although the boundary-related PMC activation patterns were consistent across internally- and externally-driven boundaries, they did not generalize across within- and between-movie boundaries. Relatedly, a recent human neurophysiological study (Zheng et al., 2022) reported that medial temporal cortex neurons distinguished within- and between-movie boundaries while subjects were watching short video clips; some neurons responded only to between-movie boundaries, whereas a separate group of neurons responded to both types of boundaries. These findings may be in line with the view that event boundaries have a hierarchical structure, with different brain areas along the information pathway reflecting different levels of boundaries, from fine-grained sensory transitions to coarse-grained situational transitions (Baldassano et al., 2017; Chang et al., 2021; Geerligs et al., 2021). However, it is still puzzling that within- and between-movie boundaries in our study produced qualitatively distinct neural patterns within a highest-order area (PMC), even though both categories consisted of prominent boundaries between situations spanning tens of seconds to several minutes. What are the crucial differences between the two levels of boundaries? One important factor might be the presence or absence of inter-event connections. Even the most salient within-movie boundaries still demand some integration of information across events, as the events are semantically or causally related, and ultimately constitute a single coherent narrative (Lee & Chen, 2021; Song, Park, et al., 2021). In contrast, an entire cluster of related events, or the narrative as a whole, might be completely “flushed” at between-movie boundaries; this difference could induce distinct cognitive states at the two levels of boundaries, giving rise to different PMC patterns.

What is the cognitive state that is generalized across internal- and external boundaries between completely different contexts, but distinct from the state evoked by boundaries within the same context? We speculate that the between-movie boundary state may be a temporary “relay” state that occurs when no one mental model wins the competition to receive full attentional focus following the flushing of the prior mental context. Namely, when one major mental context switches to another, the brain may pass through a transient off-focus (Mittner et al., 2016) or mind-blanking (Mortaheb et al., 2022; Ward & Wegner, 2013) state which is distinct from both processing external stimuli (e.g., movie watching) and engaging in internal thoughts (e.g., memory recall). This account may also explain the difference between within- vs. between-movie boundary patterns: in terms of attentional fluctuation (Jayakumar et al., 2022; Song, Finn, et al., 2021), external attention is enhanced at within-movie event boundaries (Pradhan & Kumar, 2021; Zacks et al., 2007), whereas the relay state is associated with lapses in attention (deBettencourt et al., 2018; Esterman et al., 2014). An alternative, but not mutually exclusive, possibility is that the boundary state involves the recruitment of cognitive control to resolve the competition between mental contexts. This idea is based on the observation that the areas showing relatively higher activation at between-movie boundaries overlap with the frontoparietal control network (FPCN; Vincent et al., 2008) both during encoding and recall (Figure 1B, Figure 1-figure supplement 2). As the FPCN is interdigitated with the DMN and other nearby areas within individual subjects (Braga & Buckner, 2017), relative activation of the FPCN may create the stereotyped boundary pattern in higher associative cortices. It is also noteworthy that both of these candidate cognitive states are triggered not by the onset but by the offset of a mental context; the onset would rather signal the resolution of competition between mental contexts, hence the end of those states. This dovetails with our results showing that the generalized boundary pattern appears well before movie onsets, suggesting a major contribution of offset responses.

In conclusion, we found that internally-driven boundaries between memories produce a stereotyped activation pattern in the DMN, potentially reflecting a unique cognitive state associated with the flushing and updating of mental contexts. By demonstrating stimulus-independent event segmentation during continuous and naturalistic recall, our study bridges the gap between the fields of event segmentation and spontaneous internal thoughts (also see Tseng & Poppenk, 2020). Without any task demands or external constraints, the mind constantly shifts between different internal contexts (Raffaelli et al., 2021; Sripada & Taxali, 2020). What are the characteristics of neural responses to different types of spontaneous mental context boundaries (e.g., between two different memories, between external attention and future thinking)? Is the boundary pattern observed in the current study further generalizable to mental context transitions even more stark than between-movie transitions in our experiment? Are there specific neural signatures that predict subsequent thought transitions? Future work will explore answers to these questions by employing neuroimaging methods with behavioral paradigms that explicitly and continuously track the unconstrained flow of thoughts in naturalistic settings.

## Materials and methods

Here, we provide a selective overview of procedures and analysis methods. More detailed descriptions of participants, stimuli, experimental procedures, fMRI data acquisition and preprocessing can be found in Lee & Chen (2021).

### Participants

Twenty-one subjects (12 females) between the ages of 20 and 33 participated in the study. Informed consent was obtained in accordance with procedures approved by the Princeton University Institutional Review Board. Six subjects were excluded from analyses due to excessive motion.

### Stimuli

Ten audiovisual movies (ranged 2.15 – 7.75 minutes) were used in the experiment. The movies varied in format (animation, live-action) and content. Each movie clip was prepended with a title scene where the movie title in white letters faded in and out at the center of the black screen. The movie title was shown approximately for 3 seconds of the 6-second long title scene. At the beginning of each scanning run, a 39-second long audiovisual introductory cartoon was played before the movie stimuli. The introductory cartoon was excluded from analyses.

### Experimental procedures

The experiment consisted of two phases, encoding and free spoken recall (Figure 1A), both performed inside the MRI scanner. In the encoding phase, subjects watched a series of ten short movies. Subjects were instructed to pay attention to the movies, and no behavioral responses were required. There were two scanning runs, and subjects watched five movies in each run. Stimulus presentation began 3 seconds after the first volume of each run. In the free spoken recall phase, subjects were instructed to verbally recount what they remembered from the movies, regardless of the order of presentation. Subjects were encouraged to describe their memory in their own words in as much detail as possible. A white dot was presented in the center of the black screen during the free spoken recall phase, though subjects were not required to fixate. The recall phase consisted of two scanning runs in 4 of the 15 subjects included in the analysis. The other subjects had a single scanning run. Subjects’ recall speech was audio-recorded using an MR-compatible noise-canceling microphone and then manually transcribed. The recall transcripts were also timestamped to identify the onset and offset of the description of each movie (there were no intrusions across movies during recall).

### fMRI acquisition and preprocessing

Imaging data were collected on a 3T Siemens Prisma scanner at Princeton Neuroscience Institute. Functional images were acquired using a T2*-weighted multiband accelerated echo-planar imaging sequence (TR = 1.5 s; TE = 39 ms; flip angle = 50°; acceleration factor = 4; 60 slices; 2 × 2 × 2 mm^3^). Whole-brain anatomical images and fieldmap images were also acquired. Functional images were motion-corrected and unwarped using FSL, and then coregistered to the anatomical image, resampled to the fsaverage6 cortical surface, and smoothed (FWHM 4 mm) using FreeSurfer Functional Analysis Stream. The smoothed data were also high-pass filtered (cutoff = 140 s) and z-scored within each scanning run. The first 5 volumes of encoding scanning runs and the first 3 volumes of free spoken recall scanning runs were excluded from analyses.

### Cortical parcellation and region of interest (ROI) definition

For whole-brain pattern similarity analysis, we used an atlas (Schaefer et al., 2018) which divided the cortical surface into 400 parcels (200 parcels per hemisphere) based on functional connectivity patterns (17 networks version). For region-of-interest analyses, we defined the bilateral posterior-medial cortex (PMC) by combining the parcels corresponding to the precuneus and posterior cingulate cortex within Default Network A as in our prior study (Lee & Chen, 2021). The precuneus and posterior cingulate cortex together spanned the area that showed the strongest content-and task-general boundary patterns in the whole-brain analysis (Figure 3C). The bilateral angular gyrus ROI consisted of the parcels corresponding to the inferior parietal cortex within Default Network A, B and C. The bilateral auditory cortex ROI was defined by combining the parcels corresponding to the primary and secondary auditory cortices within Somatomotor Network B.

### Univariate activation analysis

We performed whole-brain univariate activation analysis to identify brain areas that show activation changes at between-movie boundaries compared to non-boundary periods during recall (Figure 1B). The boundary periods were the first 15 seconds following the offset of each recalled movie, and the non-boundary periods were the 15 seconds in the middle of each recalled movie. Both boundary and non-boundary period time windows were shifted forward by 4.5 seconds to account for the hemodynamic response delay. We used a relatively long 15-s duration for the boundary and non-boundary periods to capture most of the boundary-related signals during recall, based on exploratory analyses that examined the time courses of univariate boundary responses (Figure 1—figure supplement 3) and boundary-triggered activation patterns (Figure 3—figure supplement 4D). For each vertex in each subject’s brain, we computed the mean boundary activation by first averaging preprocessed BOLD signals across time points within each boundary period, and then across all recalled movies. Likewise, we computed the mean non-boundary activation for each subject and vertex by first averaging preprocessed BOLD signals across time points within each non-boundary period, and then across all recalled movies. We then computed the difference between the boundary and non-boundary activation for each subject. Finally, we performed a group-level one-sample *t*-test against zero (two-tailed). The Benjamini-Hochberg procedure (False Discovery Rate *q* < .05) was applied to correct for multiple comparisons across vertices on the resulting whole-brain statistical parametric map.

### Pattern similarity analysis

We performed whole-brain pattern similarity analysis (Figure 2A) to identify brain areas that showed content-and task-general neural activation patterns associated with between-movie boundaries. For each cortical parcel of each subject’s brain, we extracted boundary and non-boundary activation patterns for each movie, separately for the encoding phase and the recall phase. Boundary patterns were generated by averaging the spatial patterns of activation within the boundary period (the first 15 seconds following the offset) of each watched or recalled movie. Non-boundary patterns were generated by averaging spatial patterns within the non-boundary period (the middle 15 seconds) of each watched or recalled movie. Again, both boundary and non-boundary time windows were shifted forward by 4.5 seconds to account for the hemodynamic response delay. We then computed Pearson correlation coefficients between the patterns within and across different movies, conditions (boundary, non-boundary), and experimental phases.

Using the resulting correlation matrix (see Figure 3A for an example) for each parcel, we first identified brain areas that showed boundary patterns which were consistent *across recalled movies* (Figure 2B). For each subject’s recall phase, we computed the mean of all pairwise between-movie correlations, separately for the boundary patterns and the non-boundary patterns. We then performed a group-level two-tailed one-sample *t*-test against zero on the mean boundary pattern correlations to test whether the boundary pattern similarity was overall positive. We also performed a group-level two-tailed paired-samples *t*-test between the mean boundary vs. non-boundary pattern correlations to test whether the boundary pattern similarity was greater than the non-boundary pattern similarity. Each of the resulting whole-brain statistical parametric maps was corrected for multiple comparisons across parcels using the Bonferroni method. Finally, we identified parcels that showed significant effects in both tests after the correction, by masking the areas that showed higher pattern similarity for the boundary than non-boundary conditions with the areas that showed overall positive similarity between boundary patterns (Figure 2B). Thus, the identified parcels showed spatially similar activation patterns across different movies at recall boundaries, and the patterns were specifically associated with boundary periods only. Likewise, we identified brain areas that showed boundary patterns which were consistent *across the encoding and recall phases* as well as across movies (Figure 2C). This was achieved by repeating the identical analysis procedures using the boundary and non-boundary pattern correlations computed across the encoding and recall phases, instead of using the correlations computed within the recall phase.

We also performed the same pattern similarity analysis in the PMC (Figure 3) and angular gyrus (Figure 3—figure supplement 2) ROIs, as done for an individual cortical parcel in the whole-brain analysis. In addition, we repeated the same analyses using shorter (4.5 seconds) boundary and non-boundary period time windows and obtained similar results (Figure 2—figure supplement 1, Figure 3—figure supplement 3).

### Comparing the onset- and offset-locked boundary patterns

To test whether the consistent activation patterns associated with between-movie boundaries were evoked by the onset or offset of a movie, we examined TR-by-TR pattern correlations across time points around the boundaries. The time points were locked to either the onset or the offset of 1) each video clip (excluding the title scene) or 2) recall of each movie. For each subject and ROI, we extracted the time series of activation patterns from 30 seconds before to 60 seconds after the onset/offset of each watched or recalled movie. We averaged the time series across movies to create a single time series of boundary-related activation patterns per experimental phase. We then computed Pearson correlation coefficients across different time points in the time series of mean activation patterns within each experimental phase (i.e., encoding-encoding and recall-recall correlation; Figure 3—figure supplement 4) or between phases (i.e., encoding-recall correlation; Figure 3D, Figure 3—figure supplement 2D). Finally, we performed two-tailed one-sample *t*-tests against zero on each cell of the time-time correlation matrices from all subjects to identify the time points at which significantly positive or negative pattern correlations appeared. Bonferroni correction was applied to correct for multiple comparisons across all cells in the time-time correlation matrix.

### Comparing the between-movie and within-movie boundary patterns

To test whether the recall activation patterns evoked by between-movie boundaries were similar to encoding activation patterns evoked by event boundaries within a movie, we identified the strongest event boundaries within each movie. We utilized the fine-grained event boundaries defined in our previous study (Lee & Chen, 2021) which divided the ten movie stimuli into 202 events excluding title scenes (mean duration = 13.5 seconds, ranged 2 – 42 seconds). We had four independent coders watch the movie stimuli and then choose which of the fine-grained event boundaries were the most important. The coders were instructed to select the boundaries such that the ten movies were divided into 60 ± 10 events excluding title scenes. Of these, 25 event boundaries were identified as important by all four coders, which resulted in 27 “coarse” events in total (ranging between one and five events per movie; mean duration = 100.9 seconds, ranged 21 – 417 seconds). To mitigate the possibility of carryover effects from the between-movie boundaries, within-movie event boundaries that occurred within the first 45 seconds of each movie clip were excluded from the analysis, leaving 15 within-movie event boundaries in total.

We first examined whether there were consistent activation patterns following the within-movie event boundaries distinct from non-boundary patterns (Figure 4—figure supplement 1). For each subject, we generated the mean PMC activation pattern for each within-movie boundary by averaging patterns from 4.5 to 19.5 seconds following the within-movie boundary during encoding. We then computed pairwise between-movie Pearson correlations across the within-movie boundary patterns, and averaged the correlations. A two-tailed one-sample *t*-test against zero was performed to test whether the similarity between the within-movie boundary patterns was overall positive. We also computed pairwise between-movie correlations across the within-movie boundary patterns and non-boundary patterns during encoding. The non-boundary pattern for each movie was generated by averaging activation patterns within the middle 15 seconds of the movie (time window shifted forward by 4.5 seconds). A two-tailed paired-samples *t*-test was performed to test whether the similarity between within-movie boundary patterns was greater than the similarity between within-movie boundary patterns and non-boundary patterns. Two of the non-boundary periods partially overlapped with the within-movie boundary periods by 13.5 seconds and 4.5 seconds, respectively, and were excluded when correlating within-movie boundary patterns and non-boundary patterns. Note that the two non-boundary periods were included in other analyses in the current study comparing between-movie boundary patterns and non-boundary patterns. However, excluding or including the two non-boundary periods did not significantly change any of the mean pairwise between-movie correlations across 1) encoding non-boundary patterns, 2) encoding non-boundary and between-movie boundary patterns, 3) encoding non-boundary and recall non-boundary patterns, and 4) encoding non-boundary and recall between-movie boundary patterns in PMC (two-tailed paired-samples *t*-tests, all *t*(14)s < 1.45, all *p*s > .17).

We next compared the template activation pattern at the within-movie event boundaries to the pattern at between-movie boundaries (Figure 4). For each subject, we generated the mean within-movie event boundary pattern of PMC by averaging activation patterns from 4.5 to 19.5 seconds following each of the 15 event boundaries during encoding. The patterns were first averaged across all time points within each boundary period time window and then across different boundaries. Likewise, the mean between-movie boundary pattern was generated by averaging all activation patterns from 4.5 to 19.5 seconds following the offset of each movie during encoding or recall. We then computed a Pearson correlation coefficient across the mean within-movie event boundary pattern and the mean encoding or recall between-movie boundary pattern. For comparison, we computed a correlation across the encoding and recall mean between-movie boundary patterns. A two-tailed one-sample *t*-test against zero was performed to test whether the group-level similarity between the two patterns was positive. We also repeated the same pattern similarity analysis using shorter (4.5 s) time windows for the boundary periods, from 4.5 to 9 seconds following the within- or between-movie boundaries (Figure 4—figure supplement 2A).

To explore the temporal unfolding of the similarity between the within- and between-movie boundary patterns, we additionally examined the between-phase TR-by-TR pattern similarity across individual time points around the boundaries (Figure 4—figure supplement 2B). For each subject, we extracted the PMC activation pattern time series from 30 seconds before to 60 seconds after 1) each within-movie event boundary during encoding and 2) the offset of each movie during recall. The time series were averaged across boundaries within each experimental phase. We then computed Pearson correlation coefficients across different time points in the activation pattern time series between the encoding and recall phases. Finally, we performed two-tailed one-sample *t*-tests against zero on each cell of the time-time correlation matrices from all subjects to identify the time points at which significantly positive or negative pattern correlations appeared. Bonferroni correction was applied to correct for multiple comparisons across cells.

### Testing the effect of visual features

Between-movie boundary periods during encoding and those during recall shared low-level visual features (i.e., mostly blank black screen). To test whether the similar visual features produced similar activation patterns at between-movie boundaries across phases, we performed a whole-brain pattern similarity analysis (Figure 2—figure supplement 2). For each subject and cortical parcel, we computed the mean boundary and non-boundary activation patterns for each movie, separately for encoding and recall. The boundary periods were defined as the first 15 seconds following the offset of each watched or recalled movie. The non-boundary periods were defined as the middle 15 seconds of each movie. Both boundary and non-boundary time windows were shifted forward by 4.5 seconds. We then computed Pearson correlations between encoding boundary patterns and recall boundary patterns across different movies, and averaged all the correlations. Likewise, we computed the average correlation between boundary patterns during encoding and non-boundary patterns during recall across different movies. A group-level two-tailed paired-samples *t*-test was performed to test whether the similarity between encoding and recall boundary patterns was greater than the similarity between encoding boundary patterns and recall non-boundary patterns, even though boundary and non-boundary patterns were visually identical during recall. The resulting whole-brain map was corrected for multiple comparisons across parcels using the Bonferroni method.

### Testing the effect of audio amplitudes

Brief periods of silence were present at transitions between movies during both encoding and recall. During encoding, the 6-second title period between movies was silent. During recall, subjects often paused speaking for several seconds between recall of different movies. We tested whether the between-movie boundary patterns were associated with the absence of sound in general, as opposed to between-movie transitions specifically.

We first compared the activation pattern associated with any silent periods within movies during encoding and the activation pattern evoked by between-movie boundaries during recall (Figure 5). To identify all periods of silence within the movies, we extracted the audio amplitudes of the movie clips (Figure 5—figure supplement 1A) by applying a Hilbert transform to the single-channel audio signals (44.1 kHz). The audio amplitudes were downsampled to match the temporal resolution of fMRI data (TR = 1.5 s), convolved with a double-gamma hemodynamic response function, and *z*-scored across time points. The periods of silence were defined as the within-movie time points (again excluding the first 45 seconds of each movie) whose audio amplitudes were equal to or lower than the mean amplitude of the time points corresponding to the silent between-movie title periods. For each subject and ROI, we averaged the activation patterns across all time points within these within-movie silent time periods to produce the mean activation pattern associated with the absence of sound. The mean pattern was then correlated with the template between-movie boundary pattern produced by averaging 4.5 – 19.5 seconds following the offset of each movie during recall. A two-tailed one-sample *t*-test was performed to compare the group-level correlation coefficients against zero.

We additionally tested whether the time course of audio amplitude was correlated with the time course of pattern similarity (Pearson correlation) between the recall phase between-movie boundary pattern and each time point of the encoding phase data (Figure 5—figure supplements 1B, 1C, 2). The time courses were generated for all time points within each movie, excluding the first 45 seconds of each movie. We first computed each subject’s Pearson correlation coefficient between the two types of time courses. We then performed a group-level one-sample *t*-test against zero (two-tailed).

## Acknowledgements

We thank Sarah DuBrow, Christopher J. Honey, and Megan T. deBettencourt for comments on earlier versions of the manuscript. We also thank Yiyuan Zhang for assisting with within-movie event boundary identification. This work was supported by the Sloan Research Fellowship (J. C.) and Google Faculty Research Award (J. C.).

## Citation diversity statement

Recent work in several fields of science has identified a bias in citation practices such that papers from women and other minority scholars are under-cited relative to the number of such papers in the field (Caplar et al., 2017; Dion et al., 2018; Dworkin et al., 2020; Maliniak et al., 2013; Mitchell et al., 2013). Here we sought to proactively consider choosing references that reflect the diversity of the field in thought, form of contribution, gender, race, ethnicity, and other factors. First, we obtained the predicted gender of the first and last author of each reference by using databases that store the probability of a first name being carried by a woman (Dworkin et al., 2020; Zhou et al., 2020). By this measure (and excluding self-citations to the first and last authors of our current paper), our references contain 14.52% woman(first)/woman(last), 16.13% man/woman, 31.26% woman/man, and 38.09% man/man. This method is limited in that a) names, pronouns, and social media profiles used to construct the databases may not, in every case, be indicative of gender identity and b) it cannot account for intersex, non-binary, or transgender people. Second, we obtained predicted racial/ethnic category of the first and last author of each reference by databases that store the probability of a first and last name being carried by an author of color (Ambekar et al., 2009; Sood & Laohaprapanon, 2018). By this measure (and excluding self-citations), our references contain 7.21% author of color (first)/author of color(last), 14.37% white author/author of color, 23.51% author of color/white author, and 54.91% white author/white author. This method is limited in that a) names, Census entries, and Wikipedia profiles used to make the predictions may not be indicative of racial/ethnic identity, and b) it cannot account for Indigenous and mixed-race authors, or those who may face differential biases due to the ambiguous racialization or ethnicization of their names. We look forward to future work that could help us to better understand how to support equitable practices in science.

## Supplementary information

**Figure 1—figure supplement 1. Changes in univariate activation at between-movie boundaries during recall (video).** The animation shows the time series of whole-brain activation maps (BOLD signals z-scored across all volumes within a scanning run) locked to the offset of the recall of each movie, from 30 seconds before to 45 seconds after the offset. Within each of the 7.5-second time windows shown as a red bar on the time axis, BOLD signals in each vertex were averaged across time points, movies, and subjects. Blue-cyan areas indicate regions with lower-than-average activation. Red-yellow areas indicate regions with higher-than-average activation.

**Figure 1—figure supplement 2. Changes in univariate activation at between-movie boundaries during encoding (video).** The animation shows the time series of whole-brain activation maps (BOLD signals z-scored across all volumes within a scanning run) locked to the offset of each movie clip during the encoding phase, from 30 seconds before to 45 seconds after the offset. Within each of the 7.5-second time windows shown as a red bar on the time axis, BOLD signals in each vertex were averaged across time points, movies, and subjects. Blue-cyan areas indicate regions with lower-than-average activation. Red-yellow areas indicate regions with higher-than-average activation.

**Figure 1—figure supplement 3.**
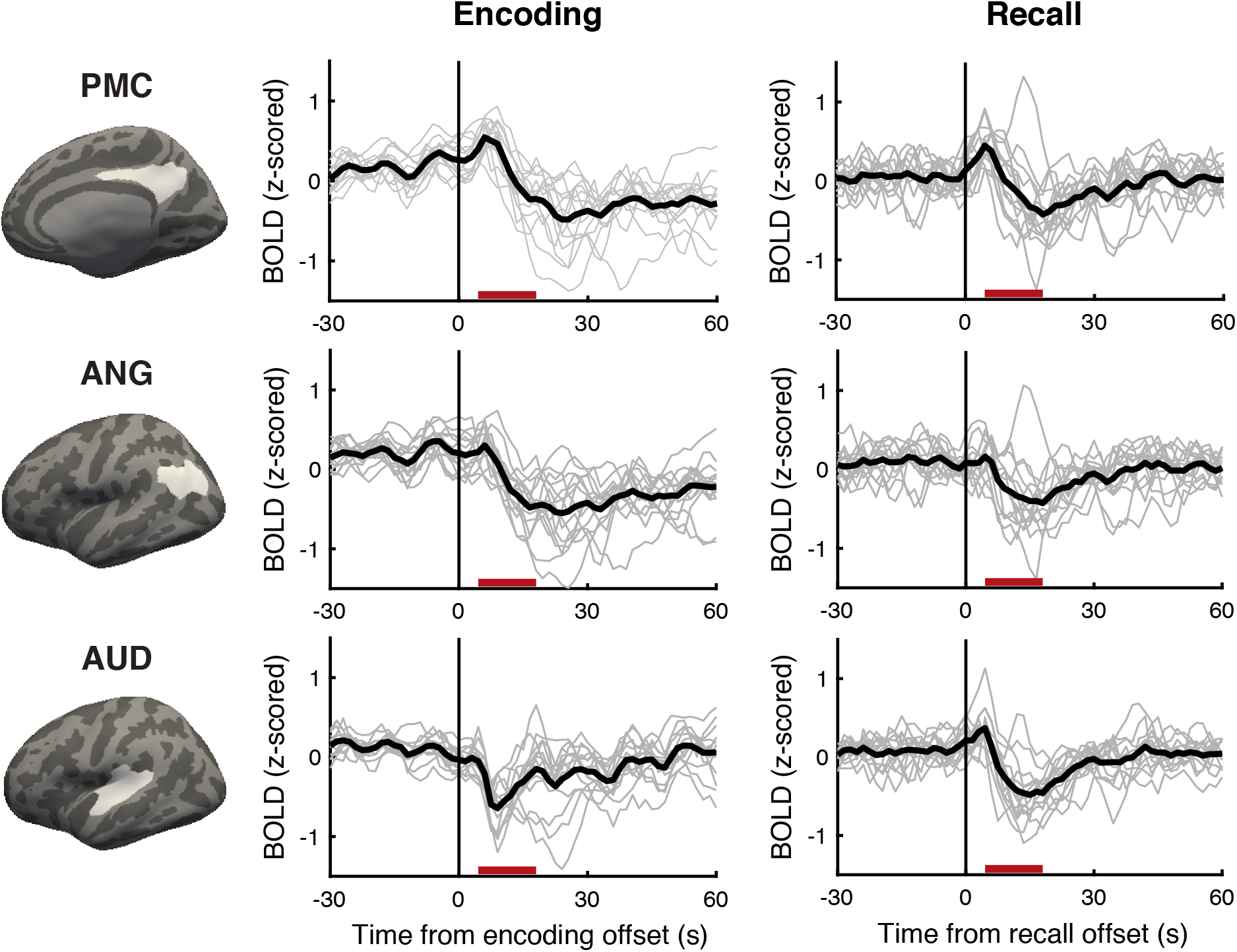
Mean activation time courses around between-movie boundaries. For each subject and region of interest (top row = PMC; middle row = ANG; bottom row = AUD), BOLD signals measured during the encoding phase (left column) or recall phase (right column) were locked to the offset of each watched or recalled movie, and then averaged across movies. Thin gray lines show individual subjects’ time courses. Thick black lines show the mean time courses averaged across all subjects. Red bars on the *x* axis indicate the 15-s boundary period time window (4.5 – 19.5 seconds from the offset of each movie) used for subsequent analyses comparing the boundary and non-boundary periods. PMC = posterior medial cortex, ANG = angular gyrus, AUD = auditory cortex.

**Figure 2—figure supplement 1.**
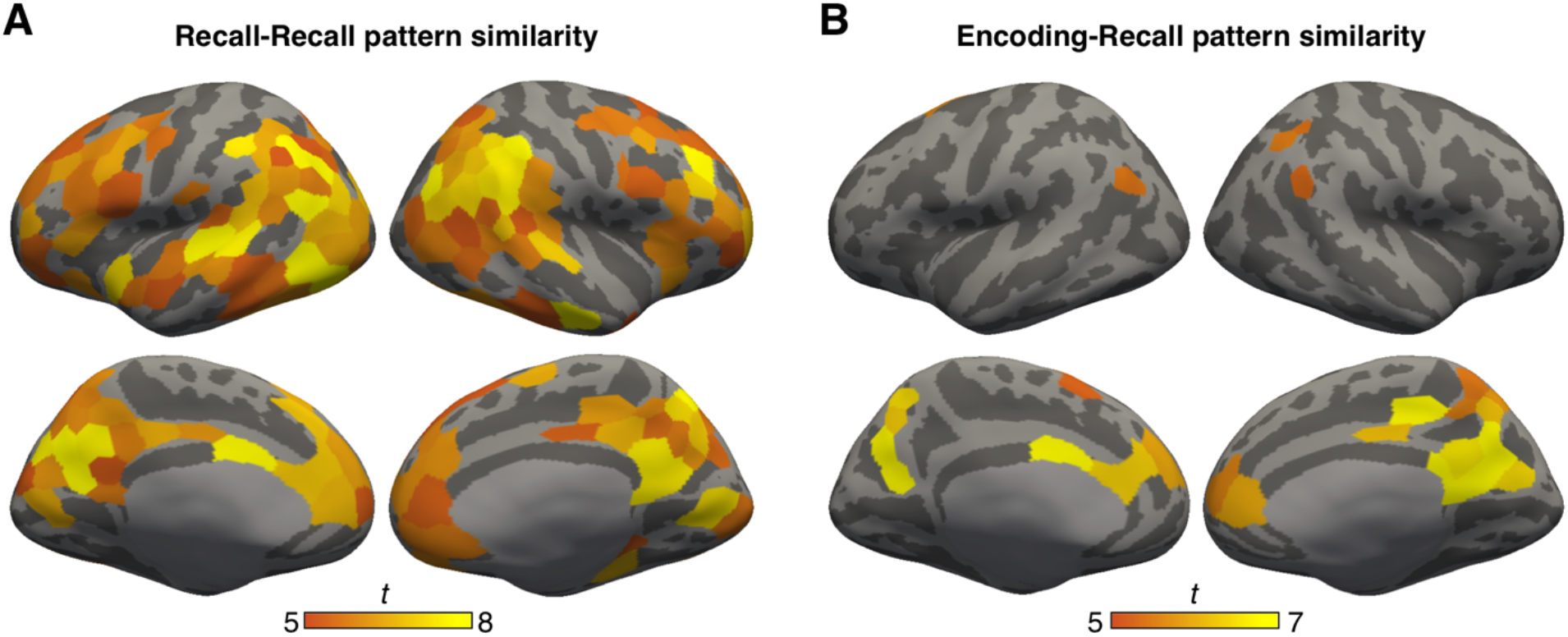
Consistent activation patterns during shorter (4.5 s) time windows following between-movie boundaries. (A) Whole-brain *t* statistic map of cortical parcels that showed consistent between-movie boundary patterns during recall. These parcels displayed significantly greater between-movie pattern similarity in the boundary condition compared to the non-boundary condition during recall. The map was masked by parcels that showed significantly positive between-movie pattern similarity in the boundary condition during recall. Both effects were Bonferroni corrected across parcels (*p* < .05). (B) Whole-brain *t* statistic map of cortical parcels that showed consistent between-movie boundary patterns across encoding and recall. These parcels displayed significantly greater between-movie and between-phase pattern similarity in the boundary condition compared to the non-boundary condition. The map was masked by parcels that showed significantly positive between-movie and between-phase pattern similarity in the boundary condition. Both effects were Bonferroni corrected across parcels (*p* < .05). For both (A) and (B), boundary periods were defined as 4.5 – 9 seconds from the offset of each movie. Non-boundary periods were defined as the middle 4.5 seconds of each movie, shifted forward by 4.5 seconds to account for hemodynamic response delay.

**Figure 2—figure supplement 2.**
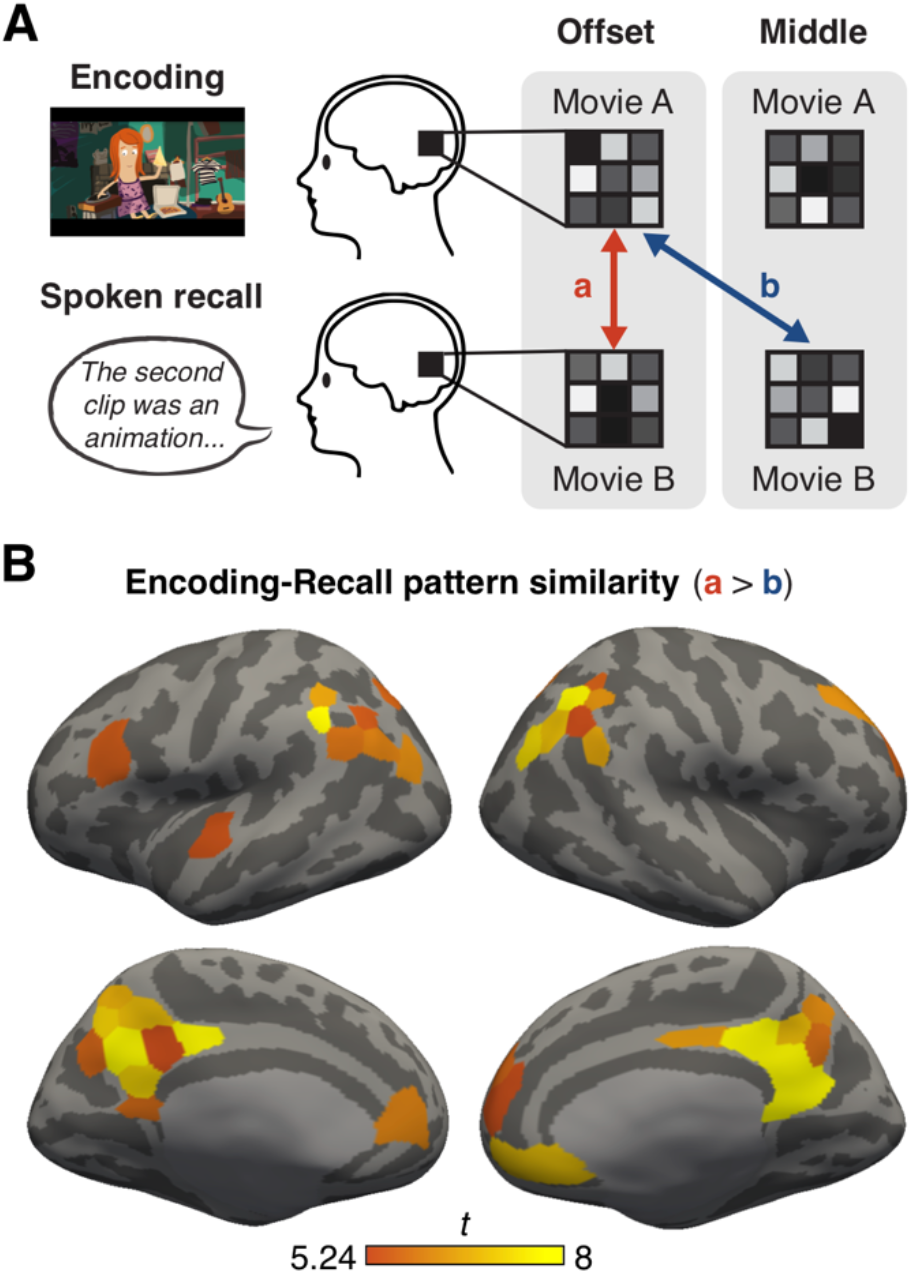
Similar visual input cannot explain between-movie boundary patterns consistent across experimental phases. (A) To test whether shared visual features (i.e., mostly-blank black screen) produced boundary (offset) patterns consistent across encoding and recall, we performed a whole-brain pattern similarity analysis. For each subject and cortical parcel, we computed the mean correlation between boundary patterns across different movies and experimental phases (a, red arrow). We also computed the mean correlation between encoding boundary patterns and recall non-boundary (middle) patterns across different movies (c, blue arrow). Note that visual input (a fixation dot) was identical across boundary and non-boundary periods during recall. The duration of boundary and non-boundary periods was 15 seconds. (B) Whole-brain *t* statistic map of cortical parcels that showed greater pattern correlations between encoding and recall boundary patterns (a, red arrow) compared to correlations between encoding boundary patterns and recall non-boundary patterns (c, blue arrow). Bonferroni correction was applied across parcels to correct for multiple comparisons (*p* < .05). Several parcels in higher associative cortices showed greater correlations between encoding and recall boundary patterns, suggesting that low-level visual features contributed little to the consistent boundary patterns in those areas.

**Figure 3—figure supplement 1.**
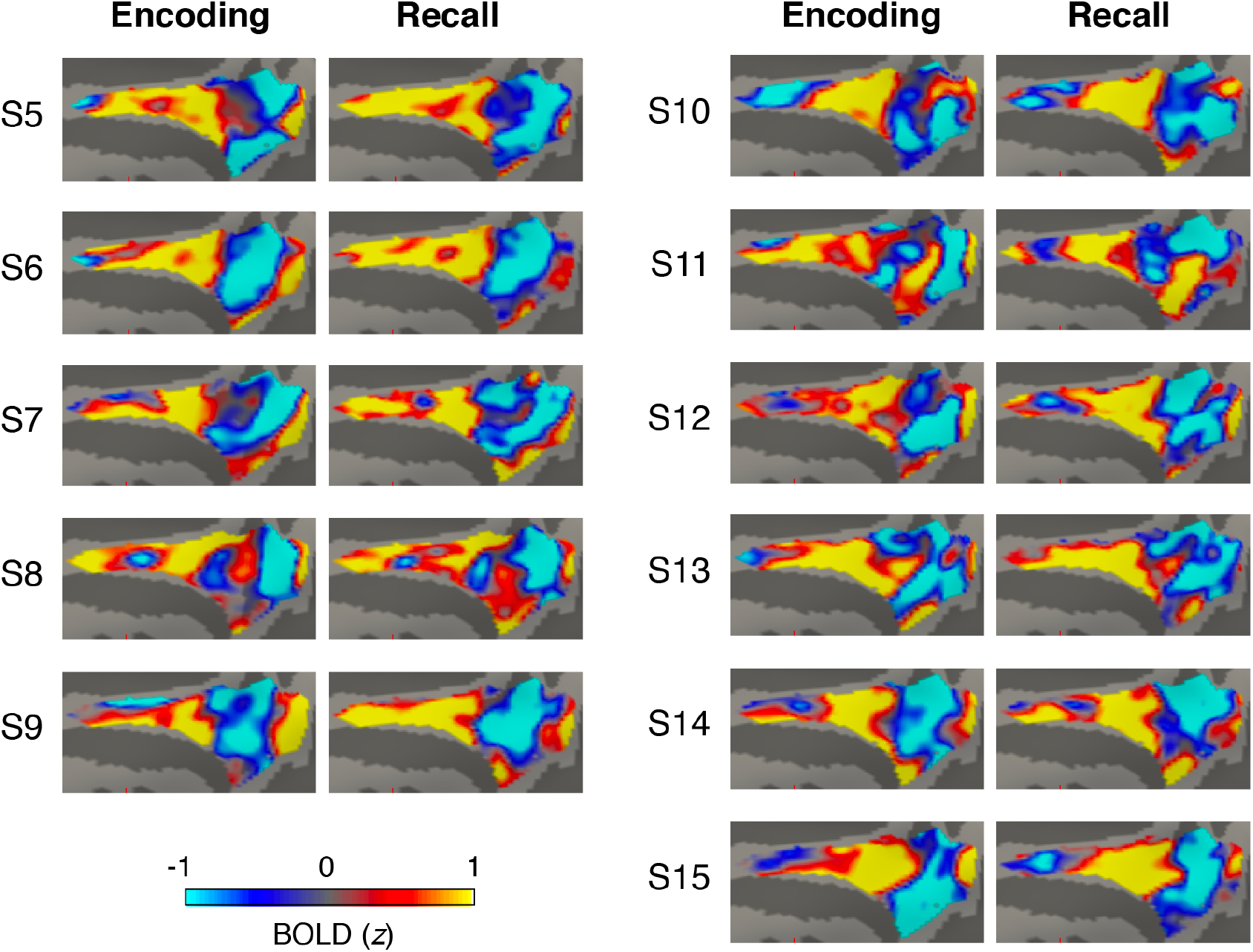
Subject-specific boundary patterns in the posterior medial cortex (PMC). The between-movie boundary patterns were averaged across all movies and then *z*-scored across vertices within the PMC ROI mask, separately for the encoding phase and the recall phase. PMC of 11 subjects (S5 – 15) are shown on the medial surface of the right hemisphere of the fsaverage6 template brain.

**Figure 3—figure supplement 2.**
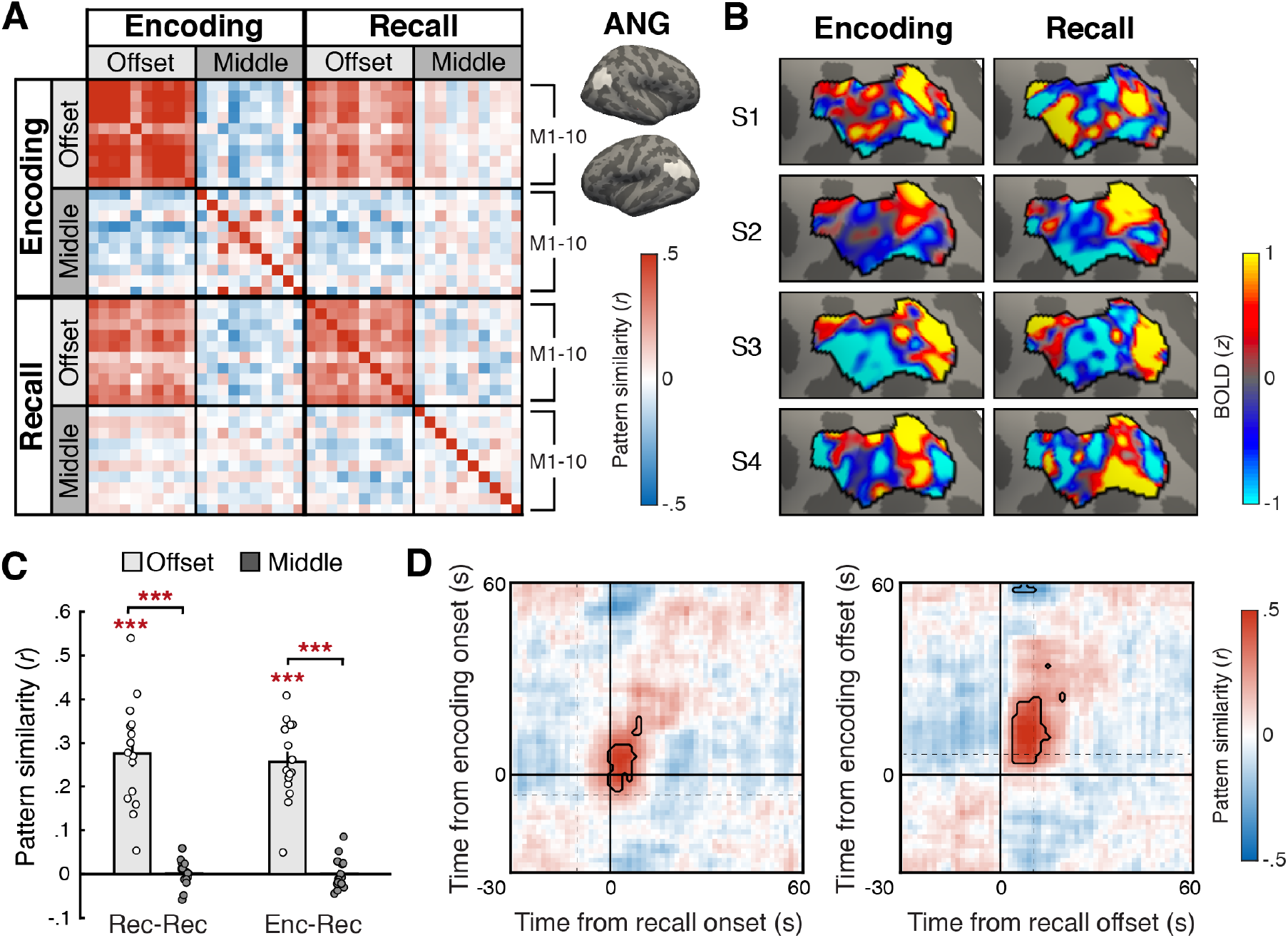
Boundary pattern in the angular gyrus (ANG). (A) ANG activation pattern similarity (Pearson correlation) between the 10 movie stimuli (M1 – 10), conditions (Offset = Boundary, Middle = Non-boundary), and experimental phases (Encoding, Recall), averaged across all subjects. The boundary pattern of a movie was defined as the mean pattern averaged across the 15-second window following the offset of the movie. The non-boundary pattern was defined as the mean pattern averaged across the 15-second window in the middle of a movie. The time windows for both boundary and non-boundary patterns were shifted forward by 4.5 seconds to account for the hemodynamic response delay. ANG regions of interest are shown as white areas on the inflated surface of a template brain. (B) Subject-specific mean activation patterns associated with between-movie boundaries during encoding (left) and recall (right). The boundary patterns were averaged across all movies and then *z*-scored across vertices within the ANG ROI mask, separately for each experimental phase. ANG (demarcated by black outlines) of four example subjects (S1 – 4) are shown on the lateral surface of the left hemisphere of the fsaverage6 template brain. (C) Within-phase (Recall-Recall) and between-phase (Encoding-Recall) pattern similarity across different movies, computed separately for the boundary (Offset) and non-boundary (Middle) patterns in ANG. Bar graphs show the mean across subjects. Circles represent individual subjects. Error bars show SEM across subjects. *\*\*\*p* < .001. (D) Time-point-by-time-point ANG pattern similarity across the encoding phase and recall phase activation patterns around between-movie boundaries, averaged across all subjects. The time series of activation patterns were locked to either the onset (left) or the offset (right) of each movie. Dotted lines on the left and right panels indicate the mean offset times of the preceding movies and the mean onset times of the following movies, respectively. Note that in this figure, zero corresponds to the true stimulus/behavior time, with no shifting for hemodynamic response delay. Areas outlined by black lines indicate correlations significantly different from zero after multiple comparisons correction (Bonferroni corrected *p* < .05).

**Figure 3—figure supplement 3.**
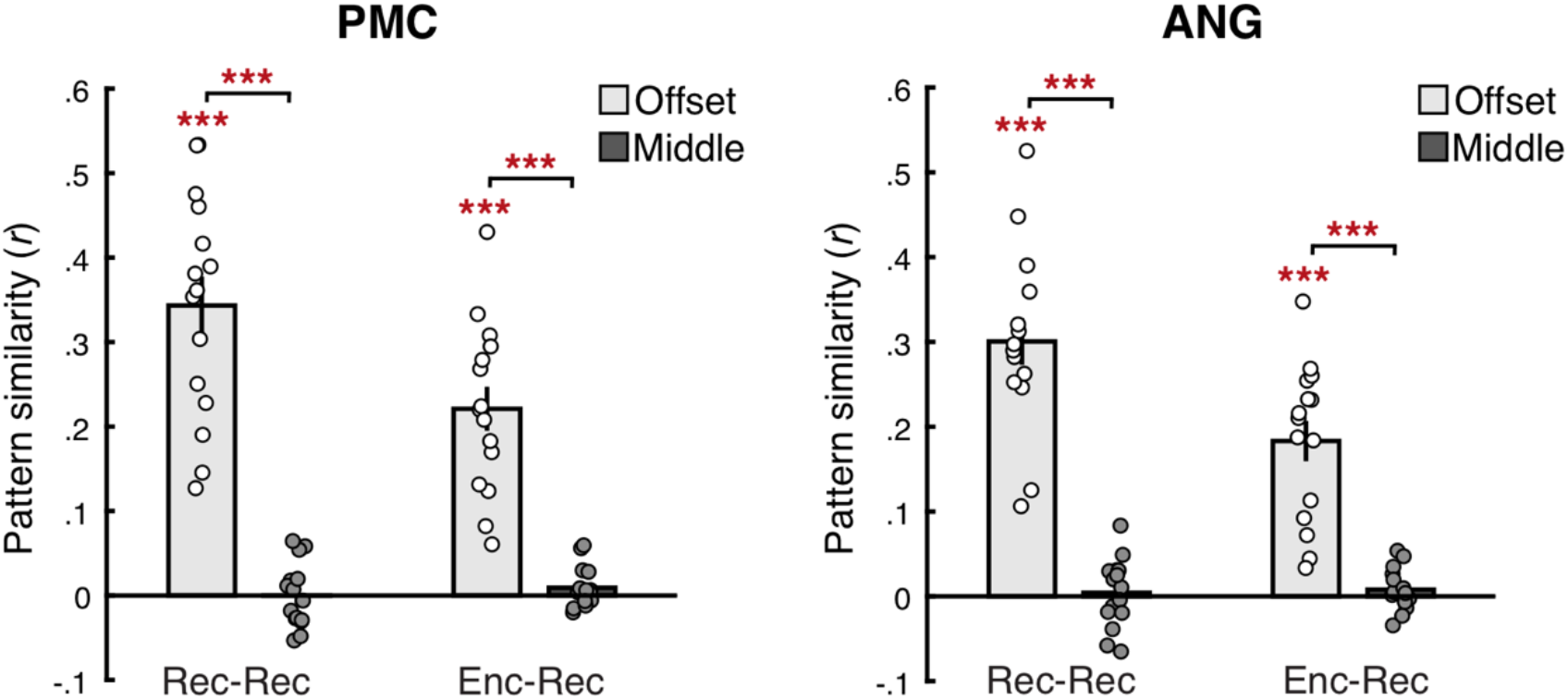
Boundary patterns in regions of interest measured during shorter (4.5 s) time windows. Within-phase (Recall-Recall) and between-phase (Encoding-Recall) pattern similarity across different movies, computed separately for the boundary (Offset) and non-boundary (Middle) patterns in the posterior medial cortex (PMC; left panel) and the angular gyrus (ANG; right panel). Boundary periods were defined as 4.5 – 9 seconds from the offset of each movie. Non-boundary periods were defined as the middle 4.5 seconds of each movie, shifted forward by 4.5 seconds to account for hemodynamic response delay. Bar graphs show the mean across subjects. Circles represent individual subjects. Error bars show SEM across subjects. \*\*\**p* < .001.

**Figure 3—figure supplement 4.**
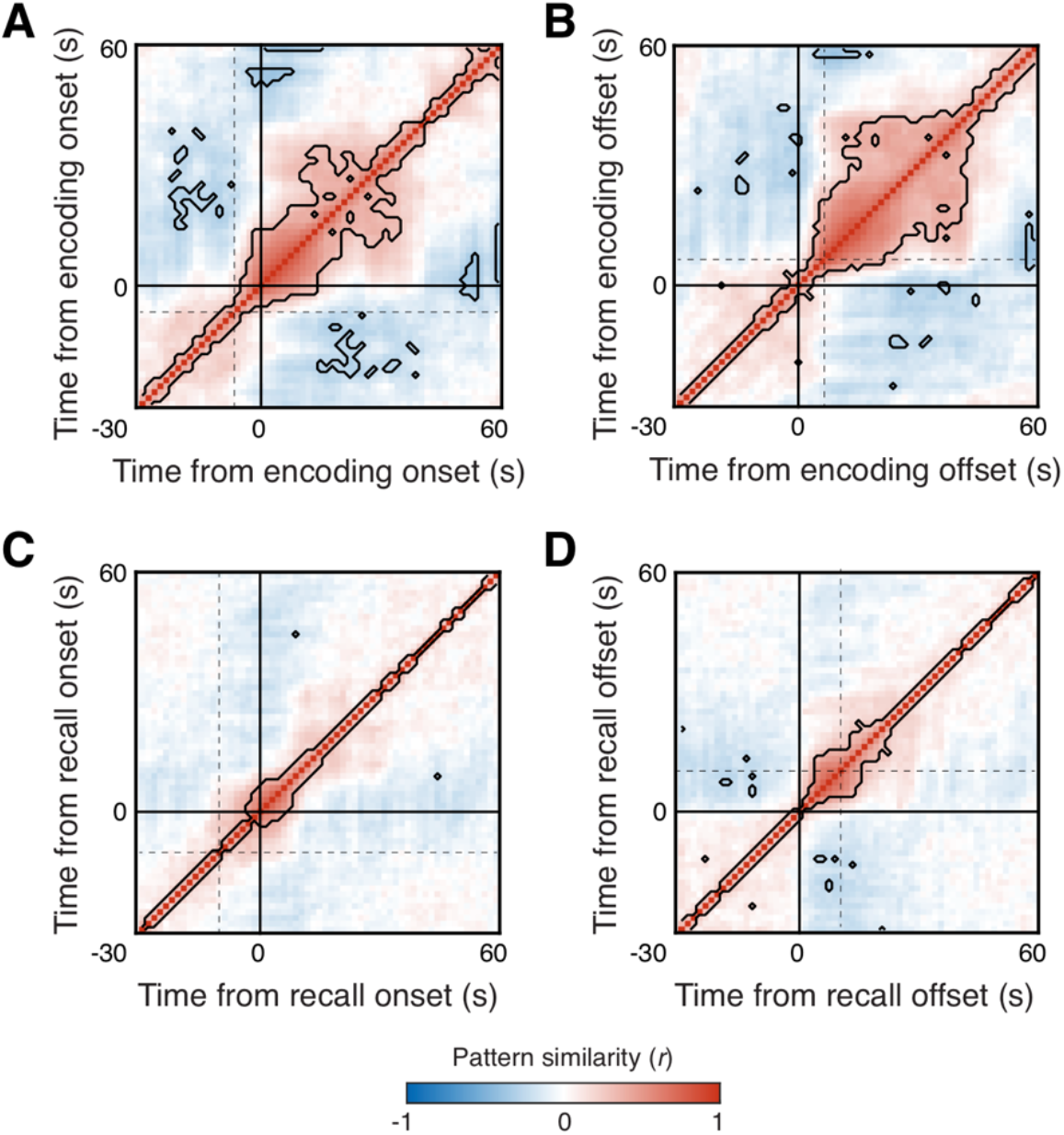
Time-time pattern similarity in the posterior medial cortex (PMC). The similarity matrices show Pearson correlations between PMC patterns across time points around between-movie boundaries during encoding (A & B) and recall (C & D), calculated within each subject and then averaged across all subjects. The time series of activation patterns were locked to either the onset (A & C) or the offset (B & D) of each movie. Dotted lines in A and C indicate the mean offset times of the preceding movies. Dotted lines in B and D indicate the mean onset times of the following movies. Note that in this figure, zero corresponds to the true stimulus/behavior time, with no shifting for hemodynamic response delay. Areas outlined by black lines indicate correlations which significantly deviate from zero after multiple comparisons correction (Bonferroni corrected *p* < .05). The boundary pattern emerged following the offsets but preceded the onsets of watched or recalled movies. In addition, the boundary pattern was stronger and lasted longer following encoding offsets compared to recall offsets; this may be because boundaries between movies were more salient during initial movie watching, as they accompanied both external and internal mental context changes whereas recall boundaries accompanied internal context changes only. Encoding boundaries were also more unpredictable and may require a more gradual build-up of the upcoming mental context, compared to self-generated boundaries between already stored memories during recall.

**Figure 4—figure supplement 1.**
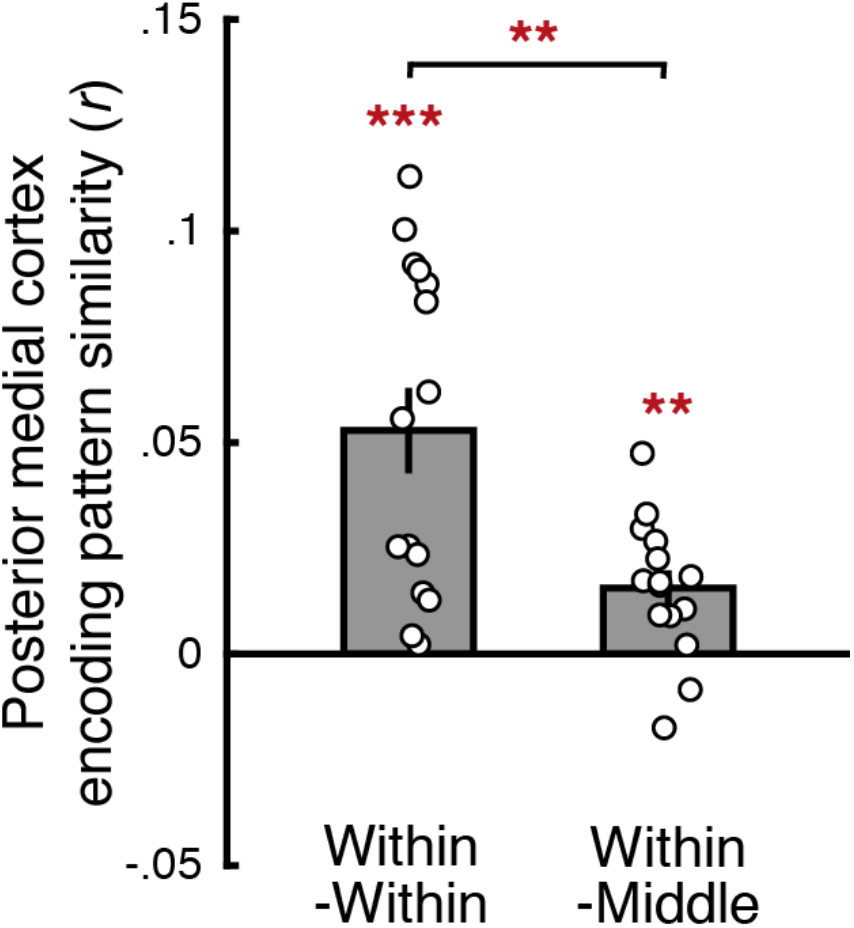
Comparing within-movie boundary patterns and non-boundary (middle) patterns in the posterior medial cortex (PMC) during encoding. The mean correlation between within-movie boundary patterns across different movies (Within-Within) was greater than zero (*t*(14) = 5.23, *p* < .001, Cohen’s *d*_z_ = 1.35, 95% CI = [.03, .07]). The mean correlation between within-movie boundary patterns and non-boundary patterns across different movies (Within-Middle) was also greater than zero (*t*(14) = 3.73, *p* = .002, Cohen’s *d*_z_ = .96, 95% CI = [.01, .02]). Critically, within-movie boundary patterns were more similar to each other than to non-boundary patterns (Within-Within vs. Within-Middle; *t*(14) = 3.29, *p* = .005, Cohen’s *d*_z_ = .85, 95% CI of the difference = [.01, .06]). The duration of within-movie boundary and non-boundary periods was 15 seconds. Two non-boundary patterns that partially overlapped with the within-movie boundary patterns were excluded from analysis. Circles represent individual subjects. Error bars show SEM across subjects. . \*\**p* < .01, \*\*\**p* < .001.

**Figure 4—figure supplement 2.**
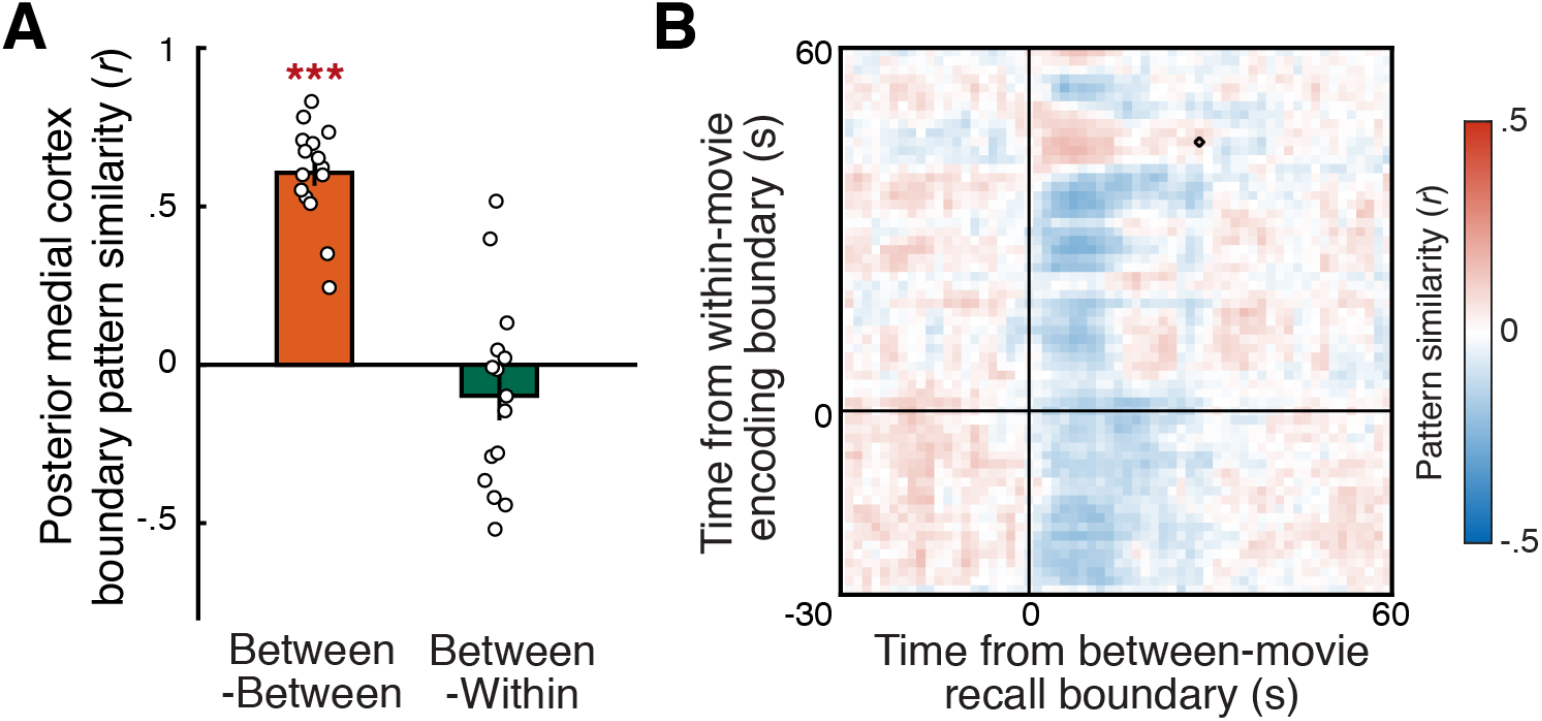
Examining the effects of boundary period time windows on the between- and within-movie boundary pattern similarity in the posterior medial cortex (PMC). (A) Pattern similarity between the template boundary patterns in PMC measured during a shorter (4.5 s) boundary period time window following the offset of each boundary. The orange bar shows the average correlation across the mean between-movie boundary patterns during encoding and recall. The green bar shows the average correlation across the mean between-movie boundary pattern during recall and the mean within-movie event boundary pattern during encoding. There was a strong positive correlation across the encoding and recall between-movie boundary patterns (*t*(14) = 15.08, *p* < .001, Cohen’s *d*_z_ = 3.89, 95% CI = [.52, .69]), whereas no such correlation was observed across the within- and between-movie boundary patterns (*t*(14) = 1.26, *p* = .23, Cohen’s *d*_z_ = .32, 95% CI = [-.26, .07]). Circles represent individual subjects. Error bars show SEM across subjects. \*\*\**p* < .001 against zero. (B) PMC pattern correlations across time points around between-movie boundaries during recall and within-movie event boundaries during encoding. The time series of activation patterns were locked to the offset of a movie or a prominent within-movie event. The correlations were first calculated within each subject and then averaged across all subjects. Time zero corresponds to the true stimulus/behavior time, with no shifting for hemodynamic response delay. Areas outlined by black lines indicate correlations which significantly deviate from zero after multiple comparisons correction (Bonferroni corrected *p* < .05). No significant positive correlations were observed across encoding and recall immediately following the within- and between-movie boundaries.

**Figure 5—figure supplement 1.**
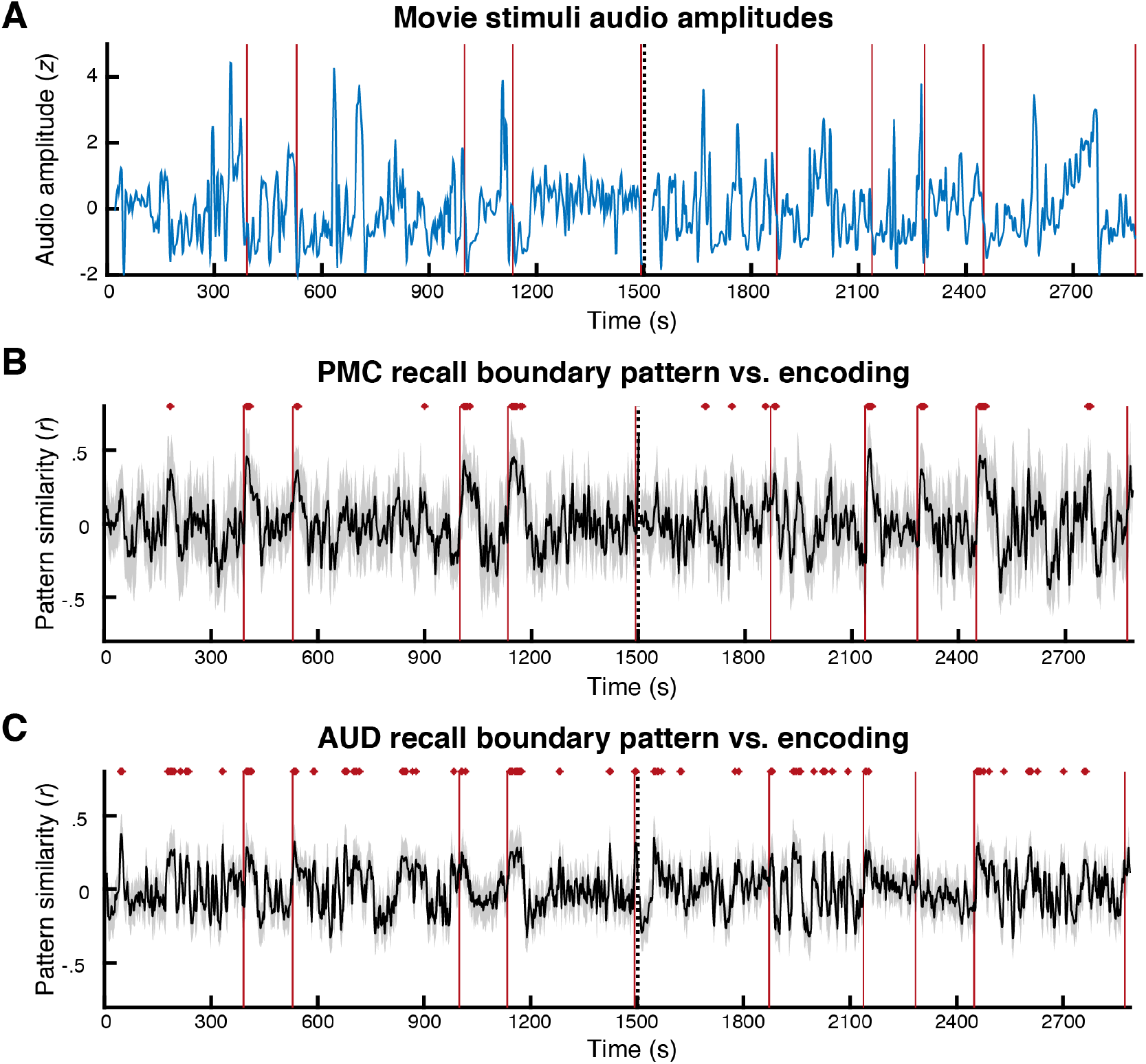
Time series of audio amplitudes during encoding and the similarity to the recall boundary pattern. (A) Audio amplitudes of the movie stimuli. The audio amplitudes were convolved with a hemodynamic response function and *z*-scored across time points. (B) Posterior medial cortex (PMC) pattern similarity (Pearson correlation) between each volume of encoding data and the template between-movie boundary pattern measured during recall (average of 4.5 – 19.5 seconds from the offset of each recalled movie). (C) Auditory cortex (AUD) pattern similarity between each volume of encoding data and the template between-movie boundary pattern measured during recall. In A, B, and C, the vertical dotted line in the middle indicates the boundary between the two encoding scanning runs. Vertical red lines indicate the offsets of each movie clip. In B and C, black lines show the mean across subjects. Shaded areas indicate the standard deviation across subjects. Red dots mark time points that showed significantly positive pattern correlations after multiple comparisons correction across time points (Bonferroni corrected *p* < . 05).

**Figure 5—figure supplement 2.**
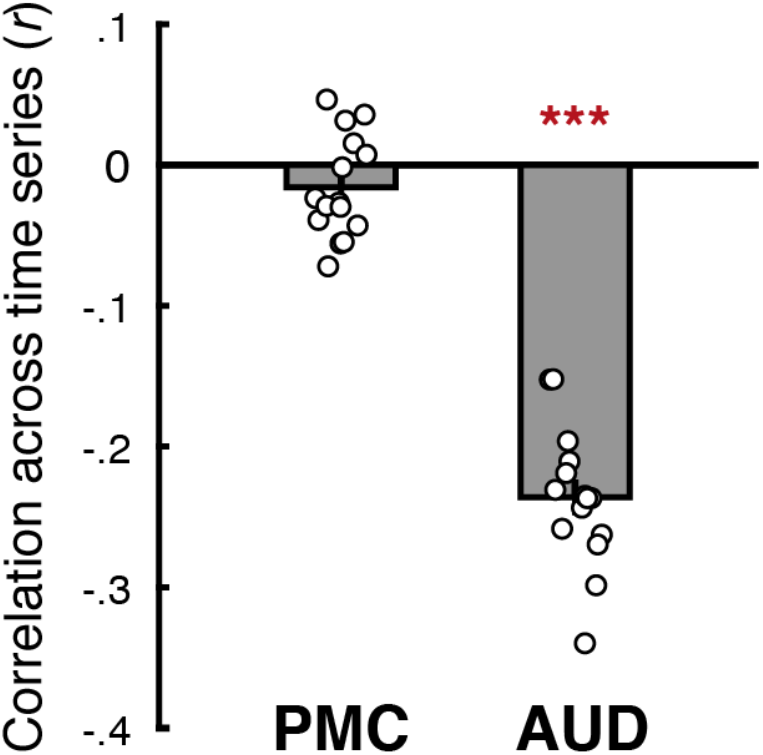
Relationship between audio amplitudes during encoding and the similarity to the recall boundary pattern. We computed correlations between the time series of audio amplitudes and the time series of similarity between the recall boundary pattern and each volume of encoding data in the posterior medial cortex (PMC) and the auditory cortex (AUD). Time points within each of the ten movies after excluding the first 45 seconds were included in the analysis. PMC correlations were not significantly different from zero (*t*(14) = 1.7, *p* = .11, Cohen’s *d*_z_ = .44, 95% CI = [-.04, .004]), whereas AUD showed significantly negative correlations (*t*(14) = 18.66, *p* < .001, Cohen’s *d*_z_ = 4.82, 95% CI = [−.26, −.21]). Bar graphs show the mean across subjects. Circles represent individual subjects. Error bars show SEM across subjects. \*\*\**p* < .001 against zero.

## References

Addis, D. R., Wong, A. T., & Schacter, D. L. (2007). Remembering the past and imagining the future: Common and distinct neural substrates during event construction and elaboration. Neuropsychologia, 45(7), 1363–1377. https://doi.org/10.1016/j.neuropsychologia.2006.10.016

Ambekar, A., Ward, C., Mohammed, J., Male, S., & Skiena, S. (2009). Name-ethnicity classification from open sources. Proceedings of the 15th ACM SIGKDD International Conference on Knowledge Discovery and Data Mining, 49–58. https://doi.org/10.1145/1557019.1557032

Andrews-Hanna, J. R., Reidler, J. S., Sepulcre, J., Poulin, R., & Buckner, R. L. (2010). Functional-anatomic fractionation of the brain’s default network. Neuron, 65(4), 550–562. https://doi.org/10.1016/j.neuron.2010.02.005

Baldassano, C., Chen, J., Zadbood, A., Pillow, J. W., Hasson, U., & Norman, K. A. (2017). Discovering event structure in continuous narrative perception and memory. Neuron, 95(3), 709–721.e5. https://doi.org/10.1016/j.neuron.2017.06.041

Ben-Yakov, A., & Dudai, Y. (2011). Constructing realistic engrams: Poststimulus activity of hippocampus and dorsal striatum predicts subsequent episodic memory. Journal of Neuroscience, 31(24), 9032–9042. https://doi.org/10.1523/JNEUROSCI.0702-11.2011

Ben-Yakov, A., Eshel, N., & Dudai, Y. (2013). Hippocampal immediate poststimulus activity in the encoding of consecutive naturalistic episodes. Journal of Experimental Psychology: General, 142(4), 1255–1263. https://doi.org/10.1037/a0033558

Ben-Yakov, A., & Henson, R. N. (2018). The hippocampal film editor: Sensitivity and specificity to event boundaries in continuous experience. Journal of Neuroscience, 38(47), 10057–10068. https://doi.org/10.1523/JNEUROSCI.0524-18.2018

Bonnici, H. M., Richter, F. R., Yazar, Y., & Simons, J. S. (2016). Multimodal feature integration in the angular gyrus during episodic and semantic retrieval. The Journal of Neuroscience, 36(20), 5462–5471. https://doi.org/10.1523/JNEUROSCI.4310-15.2016

Braga, R. M., & Buckner, R. L. (2017). Parallel interdigitated distributed networks within the individual estimated by intrinsic functional connectivity. Neuron, 95(2), 457–471.e5. https://doi.org/10.1016/j.neuron.2017.06.038

Brunec, I. K., Moscovitch, M., & Barense, M. D. (2018). Boundaries shape cognitive representations of spaces and events. Trends in Cognitive Sciences, 22(7), 637–650. https://doi.org/10.1016/j.tics.2018.03.013

Buckner, R. L., & DiNicola, L. M. (2019). The brain’s default network: Updated anatomy, physiology and evolving insights. Nature Reviews Neuroscience, 20(10), 593–608. https://doi.org/10.1038/s41583-019-0212-7

Bulkin, D. A., Sinclair, D. G., Law, L. M., & Smith, D. M. (2020). Hippocampal state transitions at the boundaries between trial epochs. Hippocampus, 30(6), 582–595. https://doi.org/10.1002/hipo.23180

Caplar, N., Tacchella, S., & Birrer, S. (2017). Quantitative evaluation of gender bias in astronomical publications from citation counts. Nature Astronomy, 1(6), 1–5. https://doi.org/10.1038/s41550-017-0141

Chang, C. H. C., Nastase, S. A., & Hasson, U. (2021). Information flow across the cortical timescales hierarchy during narrative comprehension. BioRxiv, 2021.12.01.470825. https://doi.org/10.1101/2021.12.01.470825

Chen, J., Leong, Y. C., Honey, C. J., Yong, C. H., Norman, K. A., & Hasson, U. (2017). Shared memories reveal shared structure in neural activity across individuals. Nature Neuroscience, 20(1), 115–125. https://doi.org/10.1038/nn.4450

Christoff, K., Gordon, A. M., Smallwood, J., Smith, R., & Schooler, J. W. (2009). Experience sampling during fMRI reveals default network and executive system contributions to mind wandering. Proceedings of the National Academy of Sciences, 106(21), 8719–8724. https://doi.org/10.1073/pnas.0900234106

Christoff, K., Irving, Z. C., Fox, K. C. R., Spreng, R. N., & Andrews-Hanna, J. R. (2016). Mind-wandering as spontaneous thought: A dynamic framework. Nature Reviews Neuroscience, 17(11), 718–731. https://doi.org/10.1038/nrn.2016.113

Clewett, D., DuBrow, S., & Davachi, L. (2019). Transcending time in the brain: How event memories are constructed from experience. Hippocampus, 29(3), 162–183. https://doi.org/10.1002/hipo.23074

Crittenden, B. M., Mitchell, D. J., & Duncan, J. (2015). Recruitment of the default mode network during a demanding act of executive control. ELife, 4, e06481. https://doi.org/10.7554/eLife.06481

deBettencourt, M. T., Norman, K. A., & Turk-Browne, N. B. (2018). Forgetting from lapses of sustained attention. Psychonomic Bulletin & Review, 25(2), 605–611. https://doi.org/10.3758/s13423-017-1309-5

Dion, M. L., Sumner, J. L., & Mitchell, S. M. (2018). Gendered citation patterns across political science and social science methodology fields. Political Analysis, 26(3), 312–327. https://doi.org/10.1017/pan.2018.12

DuBrow, S., & Davachi, L. (2013). The influence of context boundaries on memory for the sequential order of events. Journal of Experimental Psychology: General, 142(4), 1277–1286. https://doi.org/10.1037/a0034024

DuBrow, S., Rouhani, N., Niv, Y., & Norman, K. A. (2017). Does mental context drift or shift? Current Opinion in Behavioral Sciences, 17, 141–146. https://doi.org/10.1016/j.cobeha.2017.08.003

Dworkin, J. D., Linn, K. A., Teich, E. G., Zurn, P., Shinohara, R. T., & Bassett, D. S. (2020). The extent and drivers of gender imbalance in neuroscience reference lists. Nature Neuroscience, 23(8), 918–926. https://doi.org/10.1038/s41593-020-0658-y

Esterman, M., Rosenberg, M. D., & Noonan, S. K. (2014). Intrinsic fluctuations in sustained attention and distractor processing. Journal of Neuroscience, 34(5), 1724–1730. https://doi.org/10.1523/JNEUROSCI.2658-13.2014

Fox, M. D., Snyder, A. Z., Barch, D. M., Gusnard, D. A., & Raichle, M. E. (2005). Transient BOLD responses at block transitions. NeuroImage, 28(4), 956–966. https://doi.org/10.1016/j.neuroimage.2005.06.025

Geerligs, L., Gerven, M. van, Campbell, K. L., & Güçlü, U. (2021). A nested cortical hierarchy of neural states underlies event segmentation in the human brain. BioRxiv, 2021.02.05.429165. https://doi.org/10.1101/2021.02.05.429165

Hasselmo, M. E. (1995). Neuromodulation and cortical function: modeling the physiological basis of behavior. Behavioural Brain Research, 67(1), 1–27. https://doi.org/10.1016/0166-4328(94)00113-T

Hasson, U., Chen, J., & Honey, C. J. (2015). Hierarchical process memory: memory as an integral component of information processing. Trends in Cognitive Sciences, 19(6), 304–313.

Honey, C. J., Newman, E. L., & Schapiro, A. C. (2017). Switching between internal and external modes: A multiscale learning principle. Network Neuroscience, 1(4), 339–356. https://doi.org/10.1162/NETN_a_00024

Jayakumar, M., Balusu, C., & Aly, M. (2022). Attentional fluctuations and the temporal organization of memory. In PsyArXiv. https://doi.org/10.31234/osf.io/j32bn

Karapanagiotidis, T., Vidaurre, D., Quinn, A. J., Vatansever, D., Poerio, G. L., Turnbull, A., Ho, N. S. P., Leech, R., Bernhardt, B. C., Jefferies, E., Margulies, D. S., Nichols, T. E., Woolrich, M. W., & Smallwood, J. (2020). The psychological correlates of distinct neural states occurring during wakeful rest. Scientific Reports, 10(1), 21121. https://doi.org/10.1038/s41598-020-77336-z

Lee, H., & Chen, J. (2021). Narratives as networks: Predicting memory from the structure of naturalistic events. BioRxiv, 2021.04.24.441287. https://doi.org/10.1101/2021.04.24.441287

Maliniak, D., Powers, R., & Walter, B. F. (2013). The gender citation gap in international relations. International Organization, 67(4), 889–922. https://doi.org/10.1017/S0020818313000209

Manning, J. R., Hulbert, J. C., Williams, J., Piloto, L., Sahakyan, L., & Norman, K. A. (2016). A neural signature of contextually mediated intentional forgetting. Psychonomic Bulletin & Review, 23(5), 1534–1542. https://doi.org/10.3758/s13423-016-1024-7

Medvedeva, A., Saw, R., Silvestri, C., Sirota, M., Fuggetta, G., & Galli, G. (2021). Offset-related brain activity in the left ventrolateral prefrontal cortex promotes long-term memory formation of verbal events. Brain Stimulation, 14(3), 564–570. https://doi.org/10.1016/j.brs.2021.03.002

Mildner, J. N., & Tamir, D. I. (2019). Spontaneous thought as an unconstrained memory process. Trends in Neurosciences, 42(11), 763–777. https://doi.org/10.1016/j.tins.2019.09.001

Mitchell, S. M., Lange, S., & Brus, H. (2013). Gendered citation patterns in international relations journals. International Studies Perspectives, 14(4), 485–492. https://doi.org/10.1111/insp.12026

Mittner, M., Hawkins, G. E., Boekel, W., & Forstmann, B. U. (2016). A neural model of mind wandering. Trends in Cognitive Sciences, 20(8), 570–578. https://doi.org/10.1016/j.tics.2016.06.004

Mortaheb, S., Calster, L. V., Raimondo, F., Klados, M. A., Boulakis, P. A., Georgoula, K., Majerus, S., Ville, D. V. D., & Demertzi, A. (2022). Mind blanking is a distinct mental state linked to a recurrent brain profile of globally positive connectivity during ongoing mentation. BioRxiv, 2021.05.10.443428. https://doi.org/10.1101/2021.05.10.443428

Pettijohn, K. A., & Radvansky, G. A. (2016). Narrative event boundaries, reading times, and expectation. Memory & Cognition, 44(7), 1064–1075. https://doi.org/10.3758/s13421-016-0619-6

Pradhan, R., & Kumar, D. (2021). Event segmentation and event boundary advantage: Role of attention and postencoding processing. Journal of Experimental Psychology: General, No Pagination Specified-No Pagination Specified. https://doi.org/10.1037/xge0001155

Radvansky, G. A., & Copeland, D. E. (2006). Walking through doorways causes forgetting: Situation models and experienced space. Memory & Cognition, 34(5), 1150–1156. https://doi.org/10.3758/BF03193261

Raffaelli, Q., Mills, C., de Stefano, N.-A., Mehl, M. R., Chambers, K., Fitzgerald, S. A., Wilcox, R., Christoff, K., Andrews, E. S., Grilli, M. D., O’Connor, M.-F., & Andrews-Hanna, J. R. (2021). The think aloud paradigm reveals differences in the content, dynamics and conceptual scope of resting state thought in trait brooding. Scientific Reports, 11(1), 19362. https://doi.org/10.1038/s41598-021-98138-x

Ramanan, S., Piguet, O., & Irish, M. (2018). Rethinking the role of the angular gyrus in remembering the past and imagining the future: The contextual integration model. The Neuroscientist, 24(4), 342–352. https://doi.org/10.1177/1073858417735514

Ranganath, C., & Ritchey, M. (2012). Two cortical systems for memory-guided behaviour. Nature Reviews Neuroscience, 13(10), 713–726. https://doi.org/10.1038/nrn3338

Reagh, Z. M., Delarazan, A. I., Garber, A., & Ranganath, C. (2020). Aging alters neural activity at event boundaries in the hippocampus and Posterior Medial network. Nature Communications, 11(1), 3980. https://doi.org/10.1038/s41467-020-17713-4

Ritchey, M., & Cooper, R. A. (2020). Deconstructing the posterior medial episodic network. Trends in Cognitive Sciences, 24(6), 451–465. https://doi.org/10.1016/j.tics.2020.03.006

Schaefer, A., Kong, R., Gordon, E. M., Laumann, T. O., Zuo, X.-N., Holmes, A. J., Eickhoff, S. B., & Yeo, B. T. T. (2018). Local-global parcellation of the human cerebral cortex from intrinsic functional connectivity MRI. Cerebral Cortex, 28(9), 3095–3114. https://doi.org/10.1093/cercor/bhx179

Schapiro, A. C., Rogers, T. T., Cordova, N. I., Turk-Browne, N. B., & Botvinick, M. M. (2013). Neural representations of events arise from temporal community structure. Nature Neuroscience, 16(4), 486–492. https://doi.org/10.1038/nn.3331

Shin, Y. S., & DuBrow, S. (2021). Structuring memory through inference-based event segmentation. Topics in Cognitive Science, 13(1), 106–127. https://doi.org/10.1111/tops.12505

Smallwood, J., Bernhardt, B. C., Leech, R., Bzdok, D., Jefferies, E., & Margulies, D. S. (2021). The default mode network in cognition: a topographical perspective. Nature Reviews Neuroscience, 22(8), 503–513. https://doi.org/10.1038/s41583-021-00474-4

Smallwood, J., & Schooler, J. W. (2015). The science of mind wandering: Empirically navigating the stream of consciousness. Annual Review of Psychology, 66(1), 487–518. https://doi.org/10.1146/annurev-psych-010814-015331

Smith, V., Mitchell, D. J., & Duncan, J. (2018). Role of the default mode network in cognitive transitions. Cerebral Cortex, 28(10), 3685–3696. https://doi.org/10.1093/cercor/bhy167

Song, H., Finn, E. S., & Rosenberg, M. D. (2021). Neural signatures of attentional engagement during narratives and its consequences for event memory. Proceedings of the National Academy of Sciences, 118(33). https://doi.org/10.1073/pnas.2021905118

Song, H., Park, B., Park, H., & Shim, W. M. (2021). Cognitive and neural state dynamics of narrative comprehension. Journal of Neuroscience, 41(43), 8972–8990. https://doi.org/10.1523/JNEUROSCI.0037-21.2021

Sood, G., & Laohaprapanon, S. (2018). Predicting race and ethnicity from the sequence of characters in a name. ArXiv:1805.02109 [Stat]. http://arxiv.org/abs/1805.02109

Speer, N. K., Zacks, J. M., & Reynolds, J. R. (2007). Human brain activity time-locked to narrative event boundaries. Psychological Science, 18(5), 449–455. https://doi.org/10.1111/j.1467-9280.2007.01920.x

Sripada, C., & Taxali, A. (2020). Structure in the stream of consciousness: Evidence from a verbalized thought protocol and automated text analytic methods. Consciousness and Cognition, 85, 103007. https://doi.org/10.1016/j.concog.2020.103007

Stawarczyk, D., Bezdek, M. A., & Zacks, J. M. (2021). Event representations and predictive processing: The role of the midline default network core. Topics in Cognitive Science, 13(1), 164–186. https://doi.org/10.1111/tops.12450

Tseng, J., & Poppenk, J. (2020). Brain meta-state transitions demarcate thoughts across task contexts exposing the mental noise of trait neuroticism. Nature Communications, 11(1), 3480. https://doi.org/10.1038/s41467-020-17255-9

Vincent, J. L., Kahn, I., Snyder, A. Z., Raichle, M. E., & Buckner, R. L. (2008). Evidence for a frontoparietal control system revealed by intrinsic functional connectivity. Journal of Neurophysiology, 100(6), 3328–3342. https://doi.org/10.1152/jn.90355.2008

Ward, A. F., & Wegner, D. M. (2013). Mind-blanking: when the mind goes away. Frontiers in Psychology, 0. https://doi.org/10.3389/fpsyg.2013.00650

Wen, T., Duncan, J., & Mitchell, D. J. (2020). Hierarchical representation of multistep tasks in multiple-demand and default mode networks. Journal of Neuroscience, 40(40), 7724–7738. https://doi.org/10.1523/JNEUROSCI.0594-20.2020

Yeshurun, Y., Nguyen, M., & Hasson, U. (2021). The default mode network: where the idiosyncratic self meets the shared social world. Nature Reviews Neuroscience, 1–12. https://doi.org/10.1038/s41583-020-00420-w

Zacks, J. M. (2020). Event perception and memory. Annual Review of Psychology, 71(1), 165–191. https://doi.org/10.1146/annurev-psych-010419-051101

Zacks, J. M., Braver, T. S., Sheridan, M. A., Donaldson, D. I., Snyder, A. Z., Ollinger, J. M., Buckner, R. L., & Raichle, M. E. (2001). Human brain activity time-locked to perceptual event boundaries. Nature Neuroscience, 4(6), 651–655. https://doi.org/10.1038/88486

Zacks, J. M., Kurby, C. A., Eisenberg, M. L., & Haroutunian, N. (2011). Prediction error associated with the perceptual segmentation of naturalistic events. Journal of Cognitive Neuroscience, 23(12), 4057–4066. https://doi.org/10.1162/jocn_a_00078

Zacks, J. M., Speer, N. K., Swallow, K. M., Braver, T. S., & Reynolds, J. R. (2007). Event perception: A mind-brain perspective. Psychological Bulletin, 133(2), 273–293. https://doi.org/10.1037/0033-2909.133.2.273

Zacks, J. M., Speer, N. K., Swallow, K. M., & Maley, C. J. (2010). The brain’s cutting-room floor: Segmentation of narrative cinema. Frontiers in Human Neuroscience, 0. https://doi.org/10.3389/fnhum.2010.00168

Zheng, J., Schjetnan, A. G. P., Yebra, M., Gomes, B. A., Mosher, C. P., Kalia, S. K., Valiante, T. A., Mamelak, A. N., Kreiman, G., & Rutishauser, U. (2022). Neurons detect cognitive boundaries to structure episodic memories in humans. Nature Neuroscience, 25(3), 358–368. https://doi.org/10.1038/s41593-022-01020-w

Zhou, D., Cornblath, E. J., Stiso, J., Teich, E. G., Dworkin, J. D., Blevins, A. S., & Bassett, D. S. (2020). Gender diversity statement and code notebook v1.0. Zenodo. https://doi.org/10.5281/zenodo.3672110

